# PFN2 and NAA80 cooperate to efficiently acetylate the N-terminus of actin

**DOI:** 10.1101/2020.07.15.202630

**Authors:** Rasmus Ree, Laura Kind, Anna Kaziales, Sylvia Varland, Minglu Dai, Klaus Richter, Adrian Drazic, Thomas Arnesen

## Abstract

The actin cytoskeleton is of profound importance to cell shape, division, and intracellular force generation. Profilins bind to globular (G-)actin and regulate actin filament formation. Although profilins are well-established actin regulators, the distinct roles of the dominant profilin, profilin 1 (PFN1), versus the less abundant profilin 2 (PFN2) remain enigmatic. Here, we define a specific role for PFN2 as a stable interactor and regulator of the actin N-terminal acetyltransferase NAA80. PFN2 binding increases the intrinsic catalytic activity of NAA80. Furthermore, binding of PFN2 to NAA80 via its proline-rich loop promotes binding between the globular domains of actin and NAA80, and thus acetylation of actin. The majority of NAA80 is stably bound to PFN2, and we propose that this complex acetylates G-actin before it is incorporated into filaments. In conclusion, we reveal a functionally specific role of PFN2, and establish the modus operandi for NAA80-mediated actin N-terminal acetylation. Data are available via ProteomeXchange with identifier PXD020188.

## Introduction

The actin cytoskeleton is a highly dynamic network of actin polymers, which is involved in diverse and critical functions of every eukaryotic cell. It supplies the scaffolding for the motor force which drives cell division and intracellular transport, maintains cell shape, and powers cell migration^1^. Actin polymer dynamics are regulated by several mechanisms, including actin-binding proteins and protein modifications^2,3^ One class of these important actin regulators are profilins. They are highly abundant, cytosolic proteins of approximately 15 kDa and bind the majority of free globular (G-)actin^3^. Profilins promote actin polymerization in three ways^3^. First, they have a high affinity for actin monomers, binding at the barbed end of G-actin and actin filaments when the terminal actin monomer is ADP-bound. This increases the rate of actin-ADP barbed end dissociation from the filaments; however, when the actin is ATP-bound, profilin affinity is low and it dissociates rapidly after delivering actin-ATP to the barbed end of growing actin filaments^4^ Thus, profilins sterically block nucleation and elongation of actin filaments at the pointed end, while they promote polymerization at the rapidly growing barbed end. Secondly, they increase the rate of ADP-ATP exchange on G-actin, thereby contributing to the polymerizable pool of actin monomers available for filament formation^5^. Finally, profilins bind a range of ligands, which coordinate cytoskeletal dynamics by mediating the interaction with actin. They do so by binding polyproline motifs through a hydrophobic surface formed by N- and C-terminal helices and an internal β-sheet^6^. This allows profilin to deliver actin monomers to polyproline-containing elongation factors like formins^7^ and Ena/VASP^8–10^, effectively increasing the rate of elongation as well as preventing filament capping^10^. Profilin also acts as a weak nucleator in concert with formin^11^. In vertebrates, two major profilin isoforms, profilin 1 (PFN1) and profilin 2 (PFN2) exist, encoded by the two genes *PFN1* and *PFN2*^5^. It has been suggested that mouse PFN2 is mainly expressed in the central nervous system^12^ Human PFN2 is expressed in several nonneuronal cell lines^13,14^ PFN1 is the most abundant isoform^12,13^, as well as the most widely expressed^15,16^. Currently, the extent and importance of profilin specialization are not known. Because of alternative splicing PFN2 has two isoforms, termed PFN2a and PFN2b^17,18^. Both are 140 amino acid residues long and differ in nine residues which are located between position 109-140. PFN1, PFN2a and PFN2b are 60.7% identical at the amino acid level and share a high degree of structural similarity, with an average RMSD of 1.13 Å between the C^α^ atoms of PFN1 and PFN2^19^ PFN1 and PFN2 bind to polyproline ligands^20^ and phosphoinositides^21^ with different affinities. PFN2 has an isoform-specific role in mouse brain development, where it contributes to neuronal architecture and appropriate dendritic complexity^22^ Neuronal mouse PFN2 associates with a subset of proteins distinct from PFN1, such as components of the WAVE complex^23^. *PFN2* knockout (KO) mice have defective actin polymerization at the synapse, leading to increased synaptic excitability and increased novelty-seeking behavior^24^. Furthermore, while PFN1 promoted invasion and metastasis in a human breast cancer model, PFN2 had the opposite effect, slowing migration and lowering the metastatic potential of cancer cells^13^.

In addition to regulation by actin-binding proteins, posttranslational modifications on actin have several regulatory roles^2^. The posttranslational N-terminal acetylation (Nt-acetylation) of actin was recently revealed to be catalyzed by NAA80/NatH^25,26^, an N-terminal acetyltransferase (NAT). NATs catalyze the transfer of an acetyl group from acetyl-coenzyme A (Ac-CoA) to a protein N-terminus, be it co-or posttranslationally^27,28^. Actin Nt-acetylation is essential for maintaining cytoskeletal morphology and dynamics, as its loss in *NAA80-KO* cells results in an increased number of filopodia and lamellipodia, as well as increased cell migration^25,29^. *NAA80*-KO cells also display increased cell size, Golgi fragmentation^30^, and an increase in filamentous actin (F-actin) content compared to wild-type cells^25^. The *Drosophila melanogaster* NAA80 structure revealed how its substrate-binding site uniquely accommodates the acidic actin N-terminus^31^. Here, we show that PFN2, and not PFN1, is the main cellular interaction partner of NAA80. NAA80 selectively binds PFN2 in the absence of actin. This binding is associated with an increase in NAA80 enzymatic activity and is contingent on a polyproline stretch unique among the NAT enzymes.

## Results

### PFN2 is a specific and direct interactor of NAA80, while PFN1 does not interact

A pressing question about the novel NAT NAA80 is whether it acts alone or if its activity or localization is directed by any protein partners, as is the case for several of the previously characterized NATs^27,28^. To address this, we employed an affinity-enrichment liquid chromatography-mass spectrometry (LC/MS) approach to screen for NAA80 interactors. After expressing V5-tagged NAA80 in HeLa cells, we performed immunoprecipitation (IP), analyzing and quantifying the IPs by LC/MS and label-free quantification (LFQ). In our screen, PFN2 (log_2_ fold enrichment: 5.62, p-value: 3.6 x 10^-6^) and actin (log_2_ fold enrichment: 7.70, p-value: 4.57 x 10^-5^) were significantly enriched in the NAA80 pulldown (Fig. 1a Table S1). Both β- and γ-actin are expressed in HeLa cells, but segregating the LFQ intensities for each actin isoform by MS is challenging because most actin peptides (excluding the N-terminus) are identical^2^. We therefore refer to the combined actin protein group, recognizing that the actin intensity is made up of the total pool of actin with uncertain relative contributions from β- and γ-actin. We performed a follow-up validation screen in HAP1 cells and again, PFN2 was identified as a significant interactor of NAA80 (log_2_ fold enrichment: 2.36, p-value: 2.7 x 10^-9^). However, in this validation screen actin was not enriched by NAA80 (log_2_ fold enrichment: −0.20, p-value: 0.173) (Fig. 1b, Table S2). We observed a weak, non-significant, enrichment of PFN1 in the HeLa screen (log_2_ fold enrichment: 1.74, p-value: 0.0095), while no PFN1 enrichment was detected in the HAP1 validation screen (log_2_ fold enrichment: 0.01, p-value: 0.5325). In both co-IP experiments, we identified peptides from both PFN2a and PFN2b isoforms (Fig. S1). This confirmed that both PFN2 isoforms are present in HeLa and HAP1 cells, and that they both interact with NAA80. Given the stochastic nature of DDA proteomics^32^ and the challenge of comparing peptide intensities between peptides which are chemically different, we did not attempt to interpret the relative signal intensities of the peptides within samples to deduce which PFN2 isoform, if any, is preferred by NAA80.

**Figure 1:**
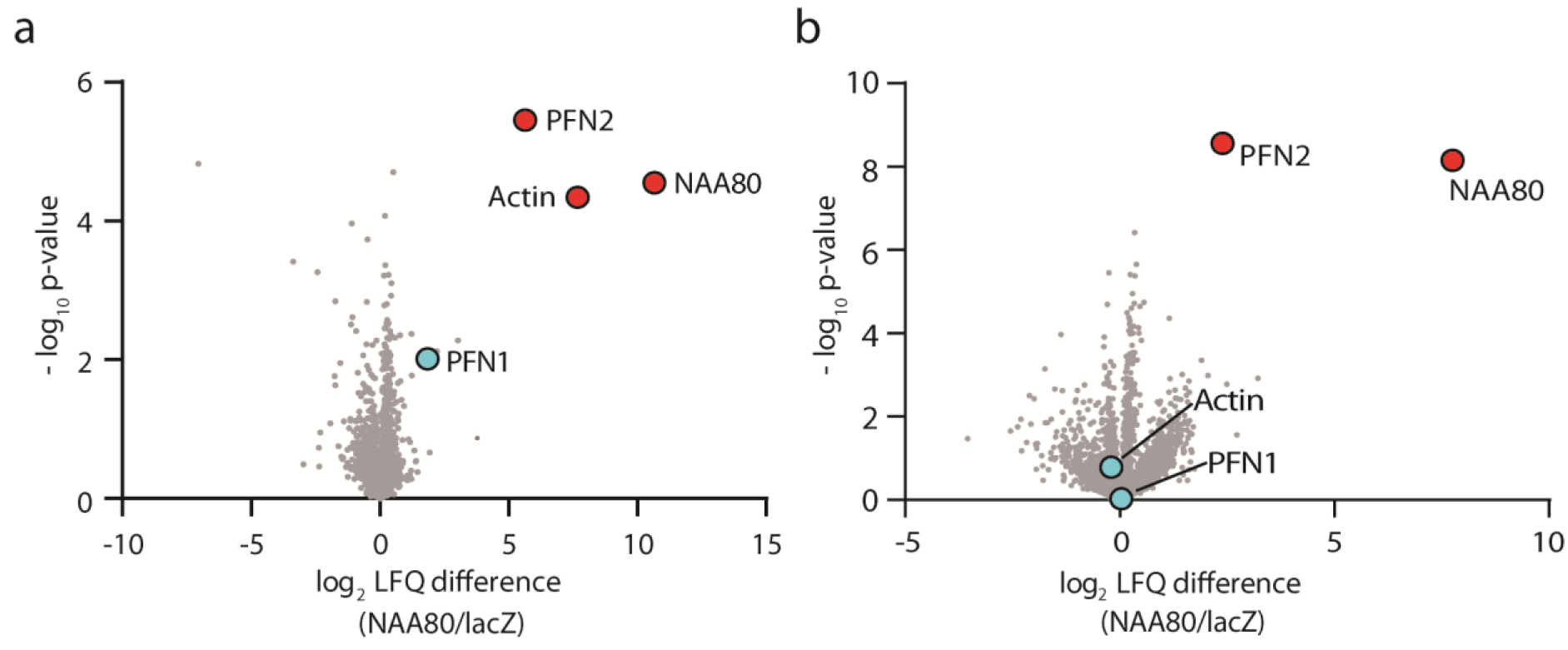
NAA80 co-immunoprecipitates PFN2. a) HeLa cells were transfected with NAA80-V5 or lacZ-V5 and b) HAP1 cells were transfected with V5-NAA80 or lacZ-V5. The V5 (IPs) were analyzed by LC/MS, and proteins were quantified by label-free quantification, comparing the intensity of each protein to the control IP (lacZ-V5) to identify interaction partners of NAA80. Red dots: significantly enriched proteins. Blue dots: selected nonenriched proteins.

While the presence of a proline-rich region (Fig. S2) of NAA80 suggested a profilin binding ability, it was unexpected to find that PFN2 and not PFN1 was enriched by NAA80. To confirm the PFN2 preference over PFN1 and the ability of NAA80 and PFN2 to interact directly and without the presence of actin, we performed analytical ultracentrifugation (AUC) experiments. First, we recombinantly expressed and purified various NAA80 constructs, PFN1, PFN2a, and PFN2b, and assessed the sample quality using several biophysical methods (Figs. S3 and S4, Table S3) before testing the effect of PFN1/2a/2b on NAA80 activity. Enzyme activity measurements (Fig. S3a-b), and circular dichroism (CD) spectroscopy and differential scanning fluorimetry (DSF) measurements (Fig. S3c, Table S3) showed that NAA80 was active and folded under the conditions used in the AUC experiments. Small-angle X-ray scattering (SAXS) measurements (Fig. S4, Table S4) and size exclusion chromatography coupled multi-angle light scattering (SEC-MALS) experiments (Fig. S4f) further confirmed a similar behavior between the profilins. All profilin isoforms were compact, monomeric, and exhibited similar molecular dimensions. Running fluorescently labeled profilins with increasing amount of full-length NAA80 allowed us to measure a potential shift in sedimentation coefficient upon NAA80 binding. Given that we identified both PFN2a and PFN2b in our LC/MS screen, we performed the AUC experiments with both isoforms. Indeed, the addition of NAA80 to either PFN2a or PFN2b resulted in a shift in the sedimentation coefficient (Fig. 2a-b). No such shift was observed with PFN1, even when we used twice the concentration as for PFN2 (Fig. 2c). Furthermore, NAA80 did not coprecipitate cellular PFN1 even in the absence of endogenous PFN2 when using *PFN2-KO* cells (Fig. 2d), indicating that endogenous PFN1 is not simply outcompeted by endogenous PFN2. Additionally, we found no further enrichment of actin by NAA80 in either of the profilin KO cells (Fig. 2d), further strengthening our conclusion that the NAA80/PFN2 interaction is actin independent and that NAA80/actin complexes are not prevalent. To determine the quantities of PFN1 and PFN2 in our HAP1 cell model we performed Western blot-based absolute quantification of PFN1 and PFN2. We have previously determined that there are approximately 22 million actin molecules per HAP1 cell, and 7000 NAA80 molecules, giving an actin:NAA80 ratio of around 3000:1^33^. Here we used the same approach, making standard curves with purified PFN1 and PFN2, and isoform-specific profilin antibodies. Our measurements indicate that there are around 4.5 million PFN1 molecules and 330,000 PFN2 molecules per cell (Fig. 2e). Thus, we report a PFN1:PFN2 ratio of 13.6:1 and PFN1:NAA80 and PFN2:NAA80 ratios of 645.0:1 and 47.5:1 respectively. The PFN1:PFN2 ratio is comparable to earlier reported measurements in non-brain tissues^12,13^. This suggests that the NAA80 preference for PFN2 cannot be explained by a mass action effect, as PFN1 is the more abundant isoform by an order of magnitude. In sum, we found that PFN2, but not PFN1, may directly bind to NAA80 *in vitro* and in cells, independent of actin.

**Figure 2:**
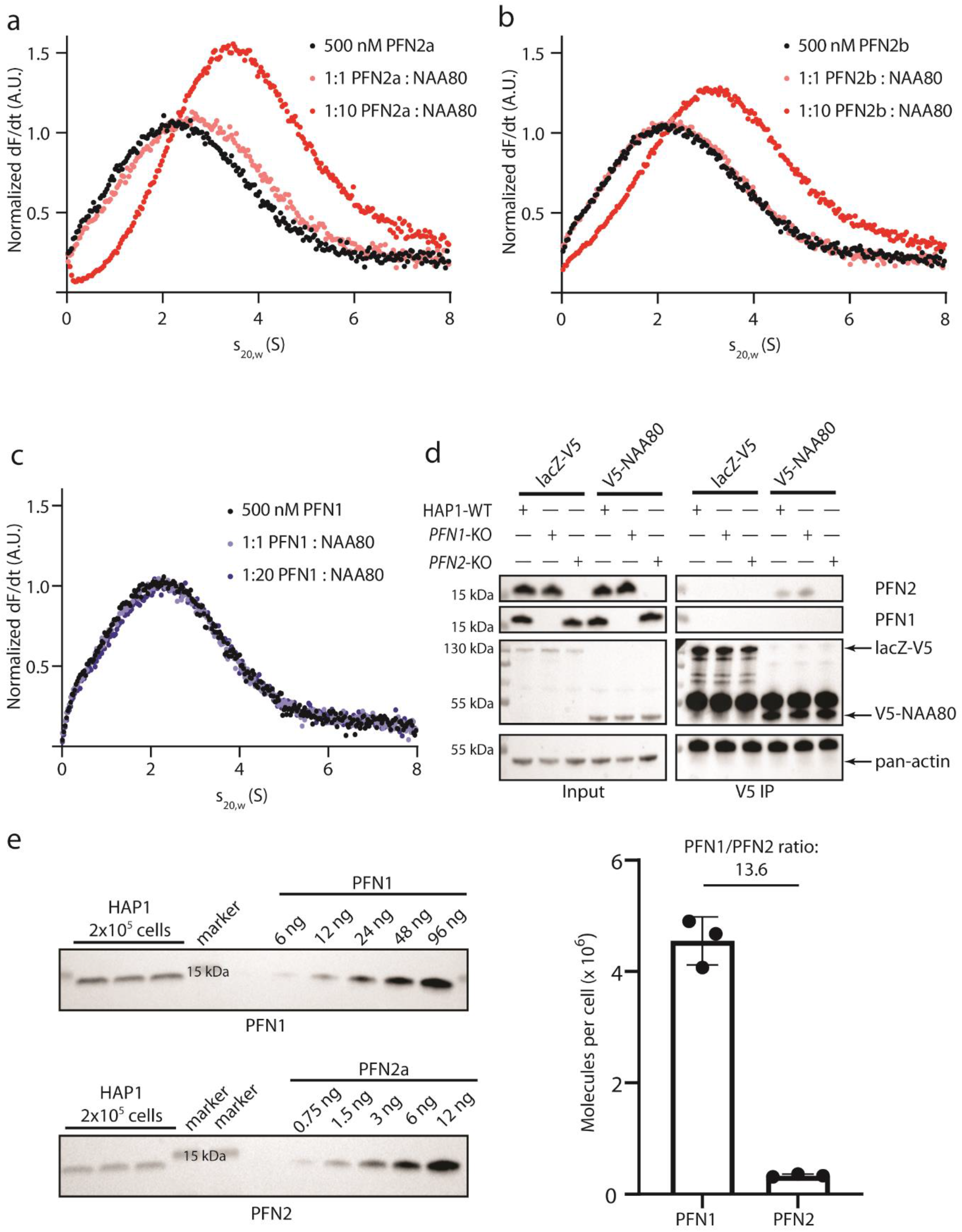
NAA80 and PFN2 specifically interact, to the exclusion of PFN1. a-c) AUC experiments with fluorescently labeled profilin and varying ratios of unlabeled NAA80. d) V5 IPs in HAP1 cells (WT, *PFN1*-KO or *PFN2*-KO) transfected with lacZ-V5 or V5-NAA80. Immunoprecipitates were probed with the indicated antibodies. Arrows indicate the expected sizes of the lacZ-V5, V5-NAA80 or actin bands. e) Western blotting-based absolute quantification of PFN1 and PFN2 in HAP1 cells, using purified proteins as standard. Right panel shows the quantification of the blots in the left panel.

### PFN2, but not PFN1, enhances the catalytic activity of NAA80 towards actin

Given that PFN2 was found to be a stable interactor of NAA80 and that profilins are established actin binding proteins, we wanted to assess whether the lack of profilins impacted global actin Nt-acetylation levels. To this end, we analyzed lysates of *PFN1-* and *PFN2-KO* cells by antibodies specific for Nt-acetylated β- or γ-actin (Fig. 3a). We found no major difference in the actin acetylation levels at steady state, and neither PFN2 nor PFN1 alone are required for actin Nt-acetylation. Next, we wanted to know whether profilin binding influences the reaction rate of NAA80. We have previously shown that preformed actin-PFN1 complexes are better substrates of NAA80 than actin alone^33^. To investigate whether NAA80-profilin is a more efficient holoenzyme than NAA80 alone, we added profilin to NAA80 before testing its activity in two different enzyme assays (Fig. 3b). In the 5,5’-dithiobis-2-nitrobenzoic acid (DTNB) assay, peptides with N-termini mimicking β-actin (DDDIA) are used as a substrate along with Ac-CoA, and the product formation is measured spectrophotometrically^34^ The DDDIA peptide is efficiently Nt-acetylated by NAA80^25^. NAA80 was preincubated with increasing molar ratios of profilin. For PFN1, we observed no increase in the NAA80 enzymatic activity. In contrast, there was a concentration-dependent increase in activity for NAA80 preincubated with PFN2a and PFN2b (Fig. 3c). NAA80 was saturated at a 5:1 PFN2:NAA80 ratio, which resulted in a 2.5-fold increase in product formation. Increasing PFN2 concentrations above this ratio did not result in any further potentiation. This suggests that PFN2 binding enhances NAA80 activity in the absence of actin, and thus without the need for the extensive NAA80-interacting interface of actin^33^, since actin is not present in this assay. Importantly, this result holds for both PFN2a and PFN2b, with a similar potentiation curve and magnitude for both isoforms. Acetylation of full-length actin can be relatively quantified by using Western blotting and antibodies specific for Nt-acetylated β- or γ-actin. We purified unacetylated β- and γ-actin from *NAA80*-KO cells^25,35^, which was used as a substrate for purified NAA80/profilin along with Ac-CoA. PFN2b significantly increased the reaction rate of NAA80 against full-length actins compared to NAA80 and actin alone, while no such effect was observed for PFN1 (Fig. 3d).

**Figure 3:**
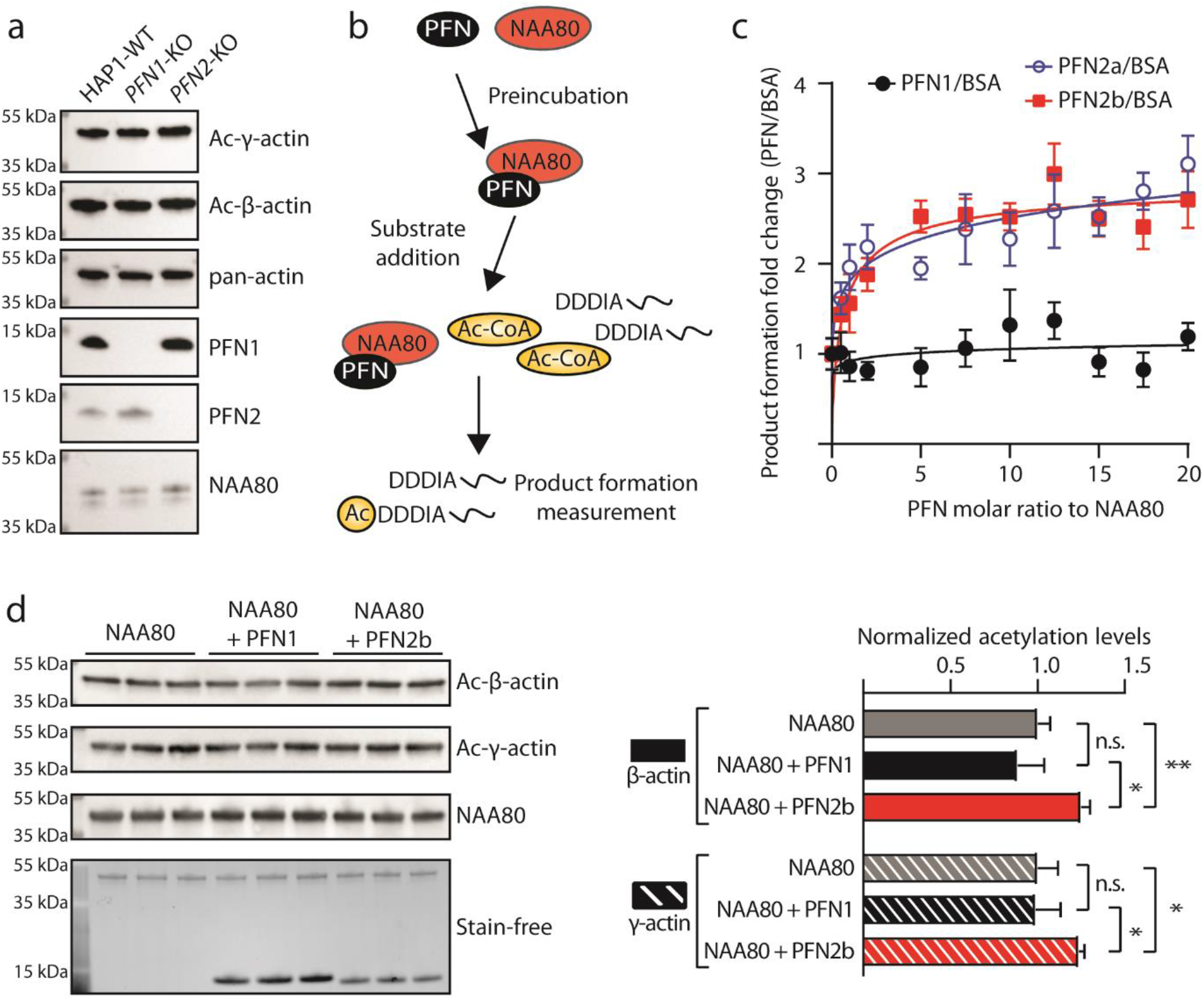
PFN2 increases the rate of actin acetylation *in vitro.* a) Actin is fully acetylated in *PFN1*-KO and *PFN2-KO* cells. Western blots of lysates from the indicated cell types. b) Schematic of the NAA80-profilin enzyme assays using β-actin N-terminal peptides (DDDIA) or full-length actin. c) NAA80/PFN DTNB enzyme assay using the N-terminal peptides of β-actin (n = 4). Enzyme reactions were run with NAA80/BSA for each ratio in parallel, and each NAA80/profilin ratio was normalized to the product formation for the corresponding NAA80/BSA ratio. Standard deviations were calculated as described in the methods section. d) *In vitro* acetylation of full-length actin in the presence of PFN1 or PFN2b. Unacetylated actin purified from *NAA80-*KO cells was used as a substrate for NAA80/profilin. Right panel shows the quantification of the blots in the left panel (n = 3 for each condition). Error bars show standard deviation. Statistical significance was determined by Student’s t-test. n.s: p > 0.05. *: p < 0.05. **: p < 0.01.

This suggested that PFN2 may act as a substrate shuttle for NAA80, using its distinct actin- and NAA80-binding domains to bring enzyme and substrate into closer proximity. However, the peptide-based DTNB assay demonstrated that actin binding is not required for this potentiation.

### NAA80 polyproline stretches are required for PFN2-NAA80 interaction and appear to be conserved among most animals

NAA80 contains several notable regions in addition to the GNAT fold, including an 82-residue N-terminus of unknown function. The N-terminus is predicted to be disordered (Fig. S5), and the presence of the initial 22 N-terminal residues distinguishes the two known NAA80 isoforms from one another. In addition, profilins are known to interact with polyproline stretches, and NAA80 contains three such regions in its β6-β7 loop, henceforth referred to as P123 (Fig. 4a). To assess whether polyprolines are a general feature of NAA80, we retrieved several NAA80 sequences from UniProtKB and compared their proline content (Fig. 5d, Fig. S2). Currently, only human^25^, murine^26^ and *Drosophila melanogaster*^31^ NAA80 have been experimentally validated regarding enzyme function. The species represent several vertebrate and invertebrate taxa. NAA80 appears to be highly conserved among mammals and the average total proline content was about ~15%. For comparison, the average proline content in human proteins is 6.2%^36^. The fish species have the highest NAA80 proline content, ranging from approximately 15% in *Danio rerio* to 20% in *Takifugu bimaculatus.* NAA80 in the crustacean species *Armadillidium vulgare* and *Portunus trituberculatus*, as well as the nematode *Caenorhabditis elegans*, have proline contents between 8-10%. Insects are the exception here – in addition to *Drosophila melanogaster,* the mosquito *Anopheles gambiae* also lacks significant proline content. We aligned the NAA80 sequences between the β6- and β7-strands, and compared the proline content of the intervening regions (Fig. 4c). Species with an extended β6-β7 loop invariably have at least one polyproline stretch. The three stretches in human NAA80 (Fig. 4a) are present in the rodents as well. The fish species have longer stretches, more numerous, or both. Insects have no NAA80 polyprolines, coinciding with their lower proline content. All of these species have two profilin genes, except for the arthropods which have one profilin each. Taken together, the evolutionary conservation of polyprolines in NAA80 suggests a conserved function. We propose that this function is NAA80-PFN2 binding to facilitate rapid actin acetylation.

**Figure 4:**
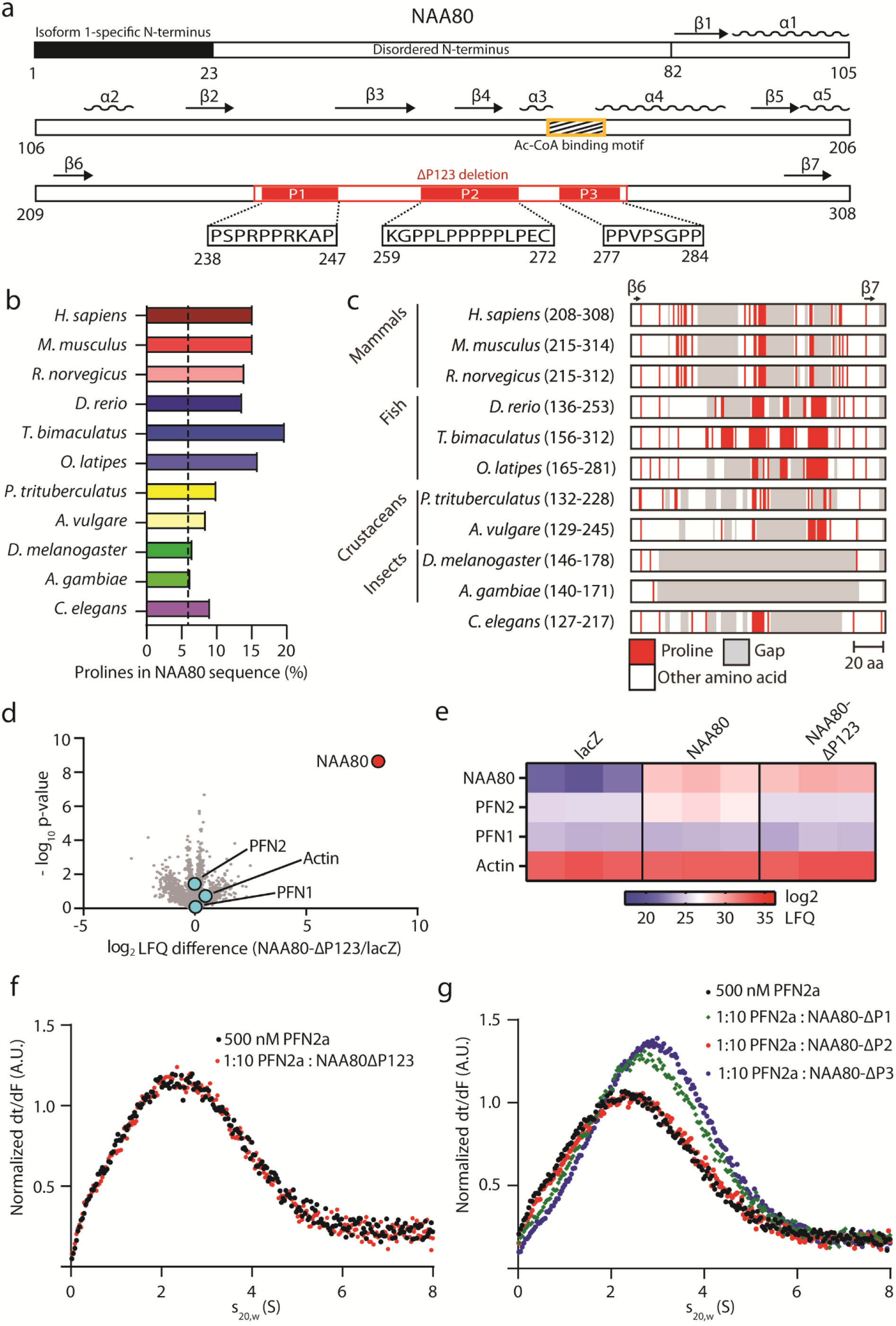
The NAA80 polyproline-rich β6-β7 loop mediates PFN2 interaction. a) Schematic of the NAA80 sequence, with secondary structure elements and regions of interest highlighted. Black: NAA80 isoform 1-specific N-terminus; yellow: Ac-CoA binding motif; red: polyproline stretches 1, 2, and 3. b) Proline content in NAA80 sequences from the indicated species: *Homo sapiens*, *Mus musculus*, *Rattus norvegicus*, *Danio rerio*, *Takifugu bimaculatus*, *Oryzias latipes*, *Portunus trituberculatus* (blue crab), *Armadillidium vulgare* (woodlouse), *Drosophila melanogaster*, *Anopheles gambiae* (mosquito), and *Caenorhabditis elegans.* Dashed line: average proline content in human genome (6.2%)^36^. c) NAA80 β6-β7 loop alignment (residue range in brackets) of NAA80 from the same species as in panel b, aligned at β6 and β7, with prolines in red and gaps in grey. d) HAP1 cells were transfected with plasmids encoding V5-tagged NAA80-ΔP123 or lacZ/β-gal. V5-tagged proteins were immunoprecipitated and interactors were identified by LC/MS and label-free quantification (LFQ), comparing the intensity of each protein to the control IP (lacZ-V5). Red dots: significantly enriched proteins. Blue dots: selected non-enriched proteins. e) Heatmap of the label-free quantification (LFQ) intensities for the indicated protein groups in each IP. f-g) AUC with labeled PFN2a and NAA80 deletion mutants: NAA80-ΔP123 (f) or NAA80-ΔP1, NAA80-ΔP2, and NAA80-ΔP3 (g).

It was of interest to define the individual contributions to PFN2 binding of the three human polyproline stretches. We performed an LC/MS interactor screen in HAP1 cells using V5-tagged human NAA80 lacking these polyproline stretches (NAA80-ΔP123) (Fig. 4d-e). Here, the PFN2 enrichment was lost in NAA80-ΔP123. We could not find any evidence of actin enrichment in this screen, nor was PFN1 enriched by NAA80-ΔP123. When we performed AUC with the same construct, it was apparent that NAA80-ΔP123 did not bind to PFN2a (Fig. 4f). We prepared deletion constructs for each stretch (NAA80-ΔP1, ΔP2 and ΔP3, lacking residues 238-247, 259-272 and 277-284, respectively), confirmed their folding and activity (Fig. S3), and performed AUC experiments (Fig. 4g). This showed that neither deletion of P1 nor of P3 had any major impact on PFN2 binding (see also Fig. S6), whereas the deletion of P2 (aa 259-272) did abolish PFN2 binding (Fig. 4g). This is in agreement with our recent study^33^, where we determined the main profilin interacting residues of NAA80 to be within the P2 stretch. Although NAA80 can also bind actin, the actin/NAA80 interaction is not essential for NAA80/profilin binding, providing further evidence that these interactions occur at different interfaces^33^.We conclude that human NAA80 binds specifically to PFN2 and that this binding is mediated through a polyproline stretch located at residues 259-272.

### PFN2-interacting prolines in NAA80 are necessary for rapid actin acetylation

We next asked; does the abrogation of PFN2 binding mean that the mutant NAA80 enzyme is unable to be potentiated by PFN2 in the same way as the full-length enzyme (Fig. 3)? We found that PFN2 is not able to increase the activity of NAA80-ΔP123, consistent with the hypothesis that the polyproline stretch contains the NAA80-PFN2 interaction interface (Fig. 5a). To exclude that the shortening of the β6-β7 loop in the deletion mutant resulted in reduced potentiation, we tested a NAA80 mutant with the critical P2 segment replaced with glycine-serine repeats (polyGS2). And comparable to the NAA80-ΔP123, the NAA80-polyGS2 mutant cannot be potentiated by PFN2 (Fig. 5b). To test whether the loss of the PFN2 interaction impairs NAA80 activity in a cellular context, we employed an actin reacetylation assay. In this assay *NAA80*-KO cells, which contain no Nt-acetylated actin^25^, were transfected with V5-tagged NAA80 variants to study how quickly each variant restored actin acetylation. We developed this assay to study actin binding to NAA80, and determined that several actin-interacting residues of NAA80 are important for rapid reacetylation^33^. We here found that NAA80-ΔP123 was significantly slower in restoring actin Nt-acetylation than the wild-type NAA80 which has retained PFN2 binding (Fig. 5c, Fig. S7). This effect was observed for both β-actin and γ-actin. Of note, NAA80-ΔP123 activity alone is higher than NAA80-WT *in vitro* (Fig. S2a). After 14 hours of transfection, NAA80-ΔP123 had restored the actin Nt-acetylation signal to 64% (β-actin) and 62% (γ-actin) compared to cells transfected with NAA80-WT, demonstrating that cellular actin acetylation kinetics are slower when NAA80 is unable to bind PFN2.

**Figure 5:**
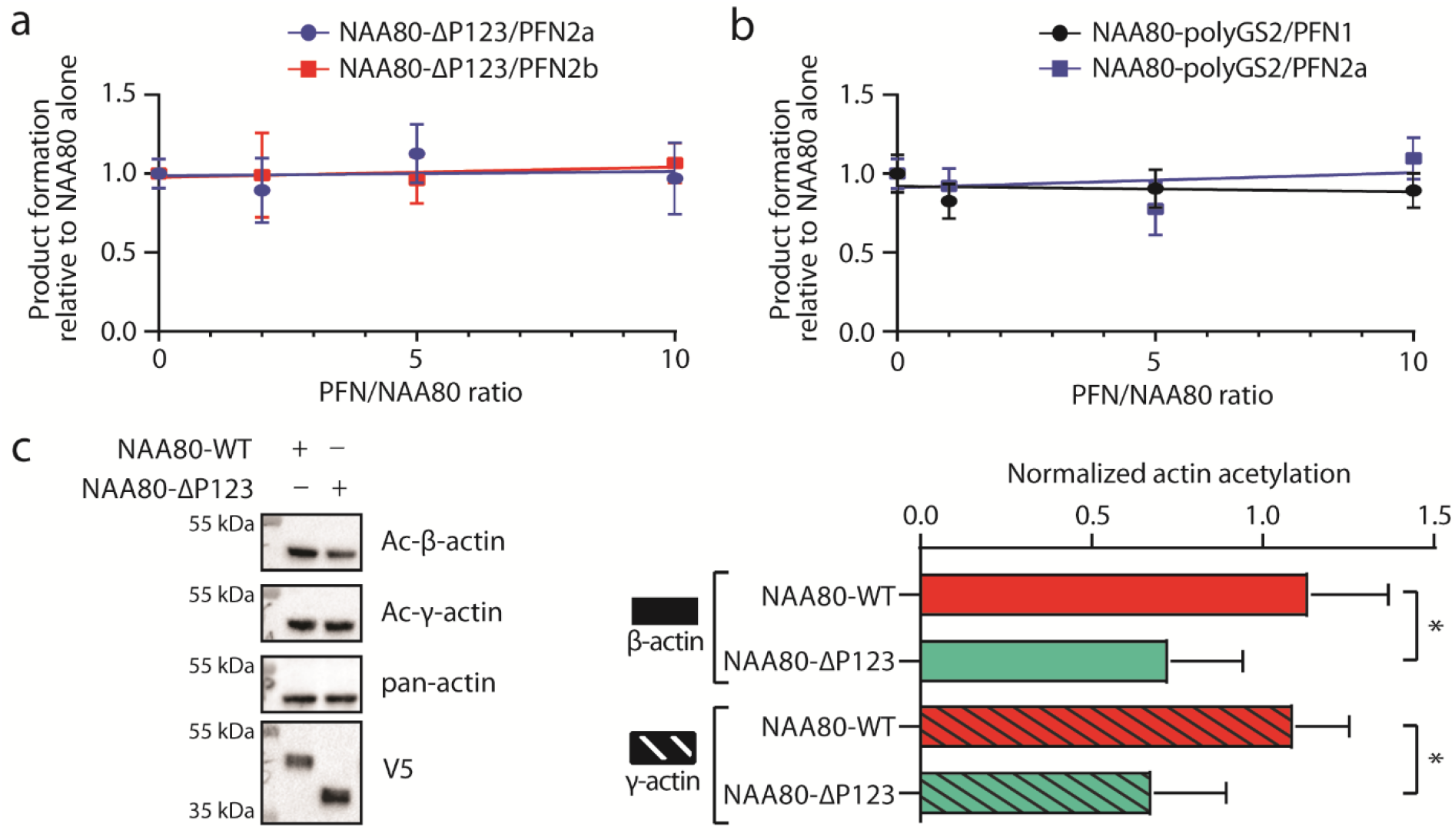
NAA80 requires PFN2 interaction for rapid actin Nt-acetylation. a) DTNB assay with NAA80-ΔP123 and varying molar ratios of PFN2a or PFN2b against DDDIA peptide (n = 3). Enzyme activity for each molar ratio was normalized to the activity of NAA80-ΔP123 alone. b) DTNB assay with NAA80-polyGS2 and varying molar ratios of PFN2a or PFN2b against DDDIA peptide (n = 3). Enzyme activity for each molar ratio was normalized to the activity of NAA80-polyGS2 alone. c) *NAA80-KO* cells were transfected with NAA80-WT or NAA80-ΔP123 for 8-14 hours before Western blotting. Left: representative blots. Right: quantification of Ac-β- and Ac-γ-actin normalized to V5 expression from independent experiments (n=4). *: p-value <0.05, as determined by two-way ANOVA. See Fig. S7 for all quantified blots.

### SAXS models of NAA80 and NAA80-ΔP123 demonstrate a high degree of conformational flexibility in the N-terminus and the β6-β7 loop

Based on our observation that the β6-β7 loop of NAA80 interacts with PFN2 (Fig. 4), we were interested in the spatial arrangement of this region in respect to the enzymatic core and its availability for interaction in solution. The crystal structure of human NAA80 was recently solved (PDB: 6NAS)^33^. However, the NAA80 construct used in the crystallization experiment was truncated at the N-terminus (ΔN-NAA80; 73 N-terminal residues missing) (Fig. 4a), and apart from a few residues which were bound to PFN1, the β6-β7 loop was not resolved^33^. The *Dm*NAA80 crystal structure also lacks the corresponding N-terminus and the extensive β6-β7 loop (PDB: 5WJD)^31^. Since these regions of NAA80 are predicted to be intrinsically disordered (Fig. S5), we performed SAXS measurements on recombinant fulllength human NAA80 (Fig. 6a, Fig. S8, Table S5). A comparison of the NAA80 scattering data to the theoretical scattering curve of the folded core, based on the *Dm*NAA80 crystal structure, demonstrated that the N-terminus and β6-β7 loop contribute to a non-globular shape of NAA80 (Fig. 6a). The NAA80 SAXS data showed that the protein has a high degree of flexibility (Fig. S8b). A SAXS model was generated by building missing loops onto a rigid body (Fig. 6b), which confirmed the extended conformation of the N-terminus and polyproline-rich β6-β7 loop. Further insight into the structural dynamics of NAA80 were gained by an ensemble optimization method (EOM) modelling approach, where we found that the N-terminus and the polyproline-rich β6-β7 loop were intrinsically disordered and could freely move in solution, without stably interacting with the globular core of the protein (Fig. 6d). The calculated *D_max_* and *R_g_* distributions of the ensemble suggested that NAA80 is exhibiting both extended and compact conformations (Fig. 6f). The rigid body and EOM models of NAA80 indicate that the N-terminus and the β6-β7 loop do not directly interact, suggesting that they have independent functions. These data demonstrate that both the N-terminus and the polyproline region are available for protein interactions in solution.

**Figure 6:**
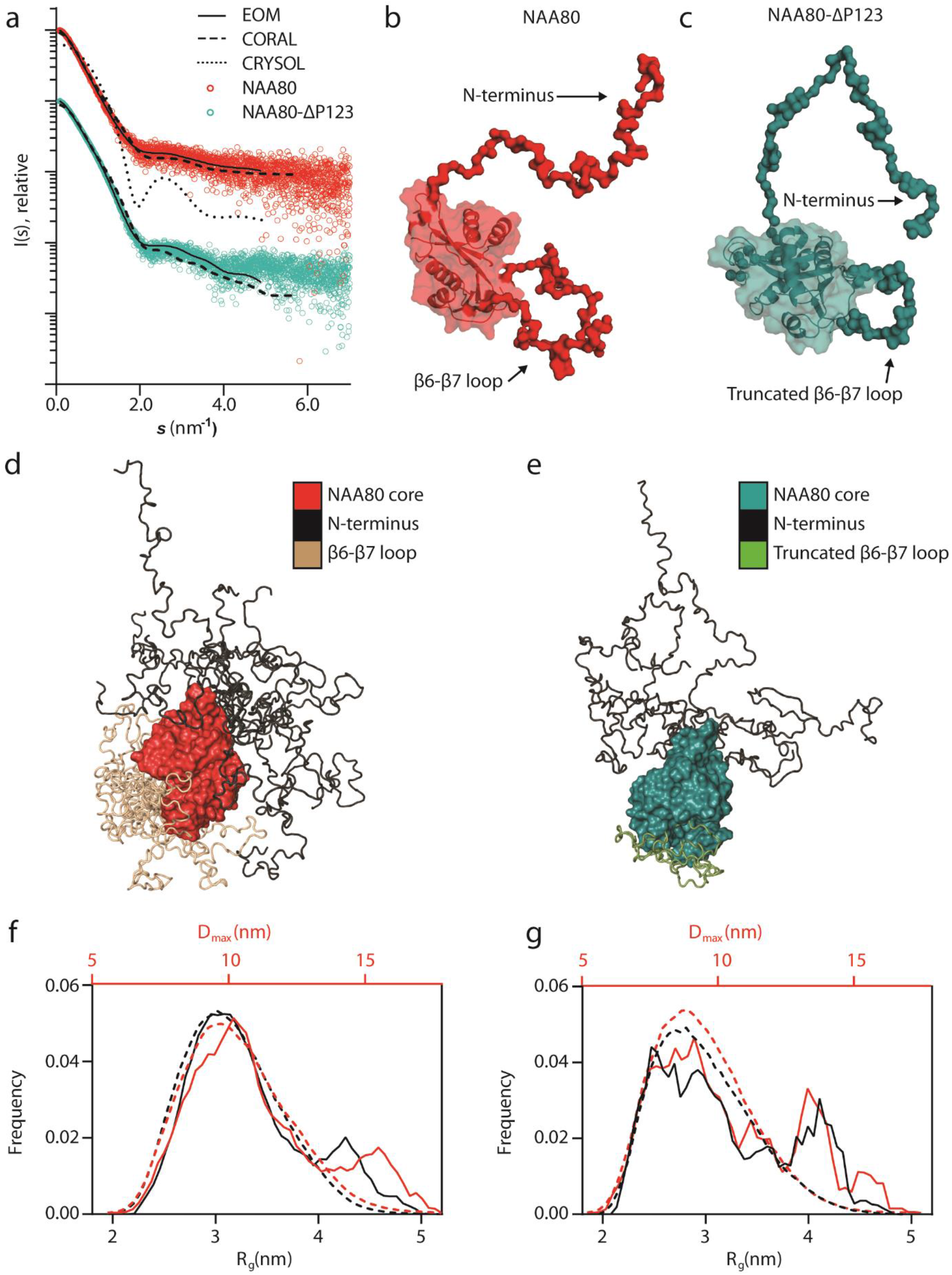
Structural information on full-length NAA80 and NAA80-ΔP123 obtained from SAXS. a) Scattering profiles of NAA80 and NAA80-ΔP123. The scattering data (dots) have been offset along the y-axis for clarity. Data fits are shown as black lines. Fit and data legend is shown as inset. The structure used for comparison by CRYSOL was *Dm*NAA80 (PDB: 5WJD,^31^) Rigid body SAXS models with missing residues modeled as dummy atoms were generated for NAA80 (b) and NAA80-ΔP123 (c) by using CORAL. d-e) EOM conformers of NAA80 (d) and NAA80-ΔP123 (e). f-g) *D_max_* (red) and *R_g_* (black) distributions from EOM analysis with ensemble frequencies (solid line) and pool frequencies (dashed lines) for NAA80 (f) and NAA80-ΔP123 (g).

We also performed SAXS on NAA80-ΔP123 to confirm the structural integrity of the polyproline deletion construct (Fig. 6, Table S5). The data verified that the GNAT fold remained intact after removing the polyproline stretch. Similarly to the full-length protein, this construct displayed a disordered N-terminus, which can move freely in solution, while the residues of the truncated β6-β7 loop are confined to a smaller conformational space (Figs. 6c, 6e, 6g). Further assessment of protein fold and thermal stability was performed using CD and DSF, respectively (Fig. S3, Table S3).

### NAA80, actin and PFN2 form a ternary complex

A structure of the ΔN-NAA80-actin-PFN1 complex was recently published^33^, showing that all three proteins make contacts with each of the other two. To determine if the cellular partner of NAA80, PFN2, can take part in similar intermolecular interactions, we performed SAXS on full-length NAA80 in complex with actin and PFN2 in solution (Fig. 7a-c, Fig. S8, Table S5). Both PFN2a and PFN2b formed ternary complexes with NAA80 and actin. CORAL models (Fig. 7b-c) were generated using the crystal structure of the ΔN-NAA80-actin-PFN1 complex (PDB: 6NAS) as rigid body. The flexible N-terminus (aa 1 – 80) and the β6-β7 loop (aa 221 – 258, 273 – 289) absent from the crystal structure were modelled based on the obtained scattering data (Fig. 7a). Our modeling confirmed that the ternary complexes formed with PFN2 have similar overall architecture as observed for PFN1. While the globular part of NAA80 forms extensive contacts in the NAA80-actin interface, the β6-β7 loop is bound to PFN2. The N-terminus of NAA80 does not seem to be involved in any of the intermolecular contacts in the NAA80-actin-PFN complex and thus may have different functions. Enrichment of PFN2, but not actin, in the IP experiments analyzed by MS (Figs. 1b and 4b-c) and Western blotting (Fig. 2d) indicated that the NAA80-PFN2 interaction is actin independent. The recently published ternary structure suggests, however, that it is primarily actin driven^33^. The capacity of NAA80 to properly acetylate actin in cells also depends on an intact actin-NAA80 interface^33^. We therefore performed AUC experiments with PFN2, NAA80 and actin to define the dependence of the NAA80-PFN2 interaction in the formation of this ternary complex relevant for cellular actin Nt-acetylation (Fig. 7 d-e). We found that while full-length NAA80 is able to form a ternary complex with PFN2-actin (Fig. 7d), NAA80-ΔP123 was not (Fig. 7e). Thus, a ternary NAA80/PFN2/actin complex may be formed and this depends on the interaction between NAA80 and PFN2.

**Figure 7:**
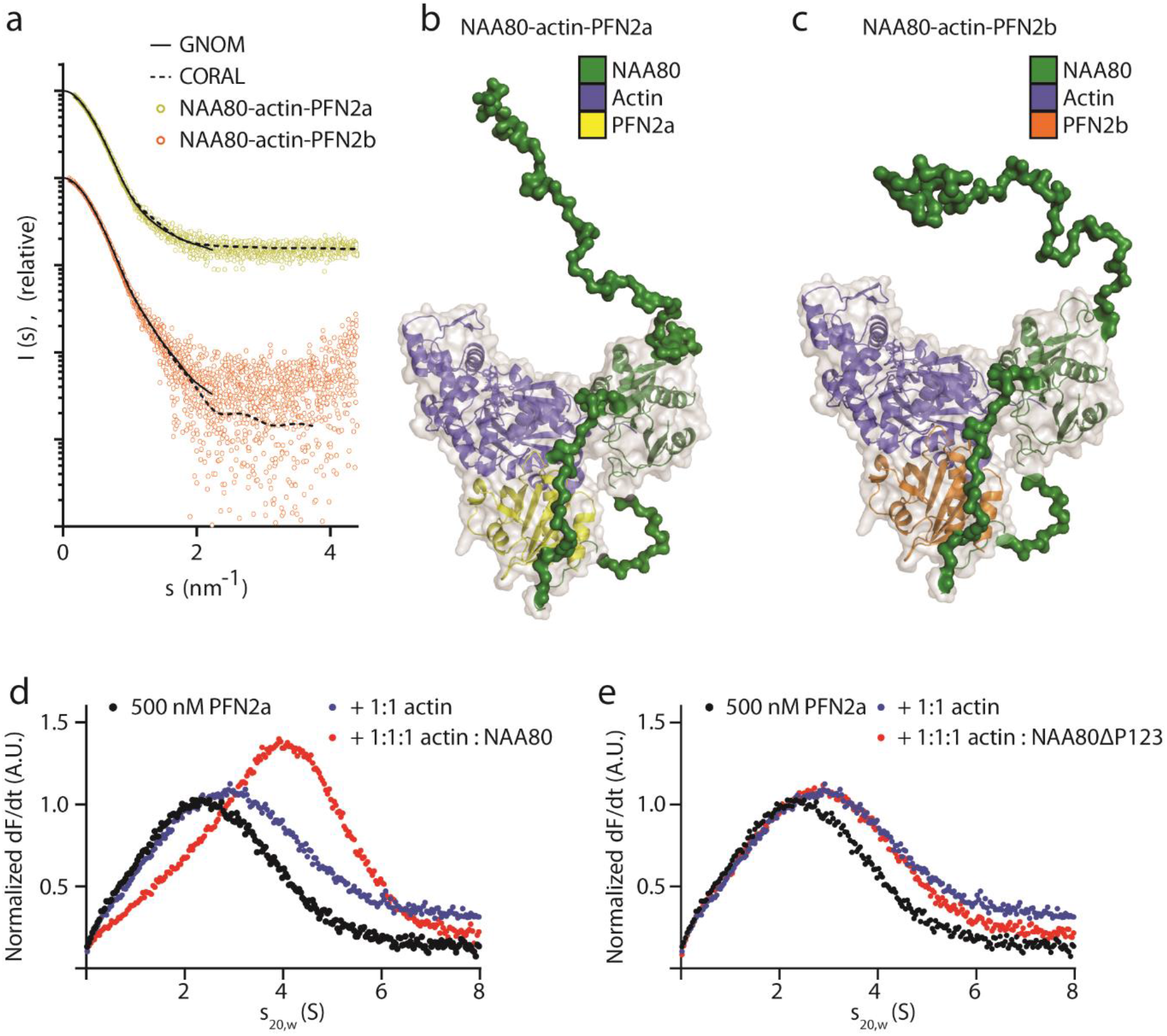
NAA80 forms a ternary complex with PFN2 and actin. a) Scattering profiles for NAA80-actin-PFN2a and NAA80-actin-PFN2b, with fits from CORAL (dashed line) and GNOM (solid line). b-c) SAXS models of NAA80-actin-PFN2a (b) or -PFN2b (c) in surface representation. The crystal structure of ΔN-NAA80-actin-PFN1 (PDB ID: 6NAS) (cartoon representation) was used as rigid body, and missing residues of the N-terminus and the β6-β7 loop were modelled by using CORAL. d-e) Analytical ultracentrifugation of labeled PFN2a, PFN2a with actin and PFN2a with actin and full-length NAA80 (d) or NAA80-ΔP123 (e).

### Cellular gel-filtration profile suggests that complexes containing actin, NAA80, and PFN2, but not PFN1, are present at low abundance

We next asked, what are the relative amounts of NAA80 in a free unbound state, bound to PFN2, or in ternary complexes with actin and PFN2? We performed gel filtration of HAP1 cell lysate and analyzed the fractions by Western blotting (Fig. 8). Gel filtration sorts proteins and protein complexes by hydrodynamic radius, which for globular proteins correlates with the molecular weight. As NAA80 (34 kDa) has a lower molecular weight than actin (42 kDa), the naïve assumption would be that actin should elute before NAA80. In addition, most cellular G-actin is profilin bound^3^, in a dimeric complex of 57 kDa. However, subjecting HAP1 lysate to gel filtration, after ultracentrifugation at 100,000 x *g* to remove F-actin, we found that NAA80 eluted before profilin-actin and indeed before the 75 kDa protein standard. The main NAA80 bands coincide with PFN2 bands, but not with PFN1 bands. This can be explained by the irregular shape of NAA80 (Fig. 6, Fig. 7), which increases its hydrodynamic radius relative to a globular protein of similar molecular weight. The presence of NAA80 and PFN2 bands in the same fractions suggest they may derive from NAA80-PFN2 complexes. However, PFN2 has other binding partners and these may account for PFN2 in these fractions. Note that this would have to be PFN2-specific binding partners, as we could not detect PFN1 in these higher molecular weight fractions.

**Figure 8:**
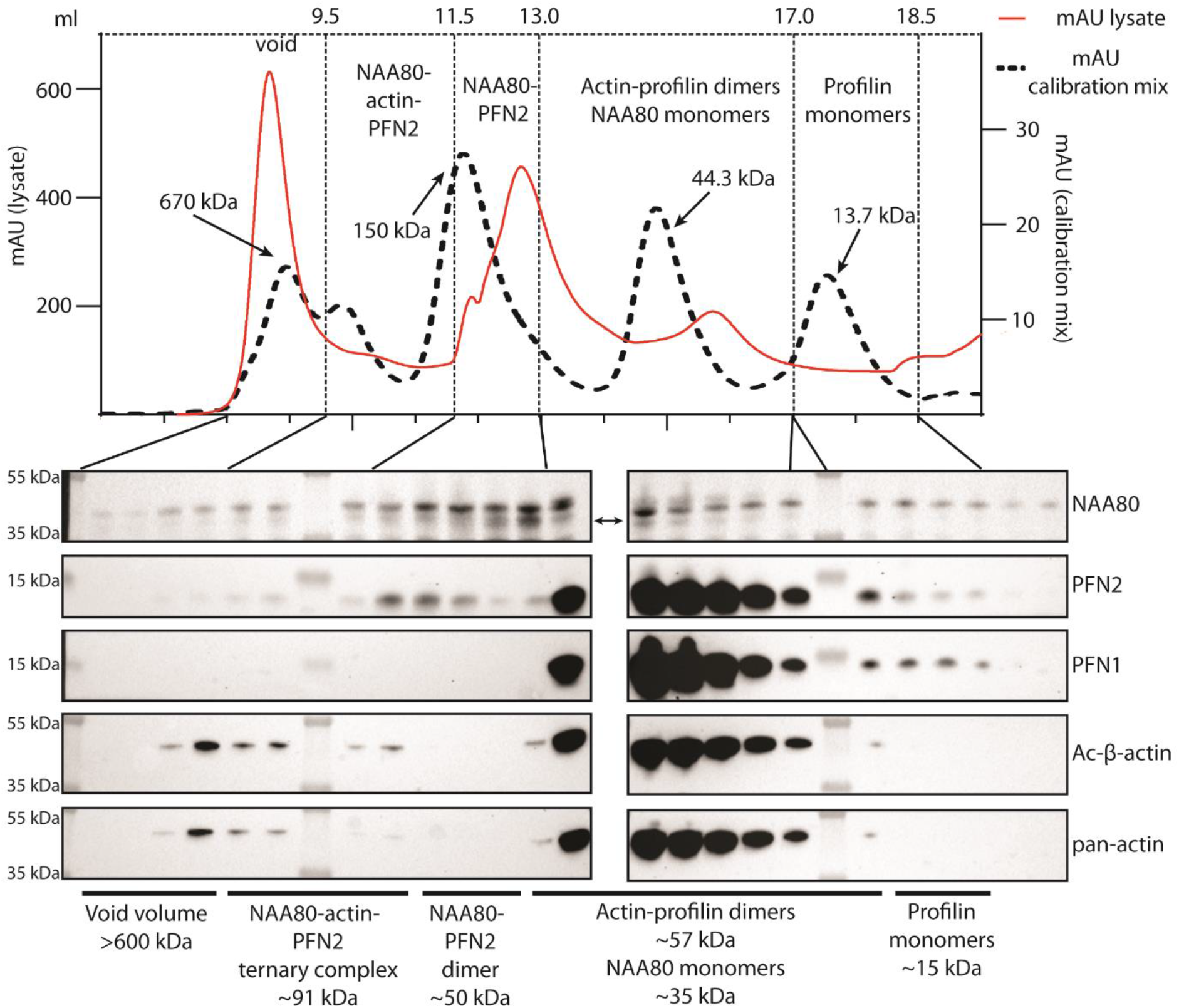
Gel filtration of cell lysates suggests the presence of low abundance NAA80-PFN2-actin complexes. Gel filtration chromatogram of HAP1 lysate and calibration mix using the same column and method. Fractions were analyzed by Western blotting and probed with the indicated antibodies. Arrows indicate sizes of standard proteins (670 kDa: bovine thyroglobulin; 150 kDa: bovine γ-globulins; 44.3 kDa: chicken egg albumin; 13.7 kDa: bovine pancreas ribonuclease A). Double-headed arrow indicates specific NAA80 band (lower).

The main fractions of profilin and actin elute between 75 and 29 kDa. These fractions are rich in profilin and actin, and presumably account for the great majority of these protein pools. Thus, we suggest that while most NAA80 is PFN2 bound, the majority of PFN2 is actin-bound and not in complex with NAA80. This is in line with the cellular ratio of PFN2 to NAA80, which we estimate to be approximately 50:1 (Fig. 2).

## Discussion

The eukaryotic NATs have different modes of operation^27^ The NatA-NatE enzymes are ribosome bound and catalyze co-translational Nt-acetylation^37,38^. Except for NatD, all of these have auxiliary subunits which anchor the enzyme complexes to the ribosome and allow catalytic subunits to Nt-acetylate nascent polypeptides^27,28,39^. For NatD, NatF and NatH it was so far believed that the enzymes NAA40^40,41^, NAA60^42,43^, and NAA80^25^ are solely responsible for their activities. Our current data reveal that NAA80 interacts with PFN2 in human cells (Figs. 1 and 2), and further that PFN2 both increases the intrinsic catalytic NAT activity of NAA80 (Fig. 3) as well as facilitating contact between NAA80 and its substrate actin (Fig. 7). Loss of the PFN2-interaction domain of NAA80 leads to inefficient actin acetylation in *NAA80-KO* cells (Fig. 5c). PFN2 may thus be considered an auxiliary subunit of NAA80 in a NatH complex. The above data are all in favor of PFN2’s candidacy as a NAT subunit; however, two other arguments should be considered. One is the apparently normal steady state level of Nt-acetylated actin in *PFN2-KO* cells (Fig. 3a). The other is the fact that there are almost 50 PFN2 molecules for each NAA80 molecule (Fig. 2e) and consequently a large fraction of cellular PFN2 is not complexed with NAA80 (Fig. 8).

Using interaction proteomics, analytical ultracentrifugation and NAA80 activity assays, we show that PFN2 is the only cellular protein which interacts stably and specifically with NAA80. The PFN1:PFN2 ratio of more than 10:1 shows that this is not due to a mass action effect, and IPs in *PFN2*-KO cells shows that even when PFN2 is absent PFN1 is unable to take its place. The present study is not the first to identify differing binding partners to the different profilin isoforms. Native PFN1 and PFN2 complexes from mouse brain were found to contain both exclusive and shared binding partners^23^. The BioPlex interactomics database^44^, as of April 2020, lists both PFN1 and PFN2 as NAA80 interaction partners in HEK293T cells. PFN1 was identified as a bait of NAA80, with NAA80 as its only currently annotated interaction partner in that database. At the time of writing, only the NAA80 IP was listed in BioPlex, not the reverse PFN1 IP. PFN2 is annotated as a NAA80 prey and *vice versa.* PFN2 has, besides actin, other interactors in this dataset, mainly actin cytoskeleton remodelers. NAA80, in addition to the profilins, are annotated as binding β-, α-cardiac and α-aortic smooth muscle actins, the transcriptional regulator BTB and CNC homolog 2 (BACH2), betaactin-like protein 2 (ACTBL2) and putative beta-actin-like protein 3 (POTEKP). Our current results indicate that PFN2 is the only reliable interactor of NAA80 in HAP1 cells, but we cannot exclude the possibility that there is context- or cell type-specific binding to PFN1 or to other actin-binding proteins. IP coupled to MS or Western blotting cannot distinguish direct from indirect binding so the possibility that some of these actin-binding proteins were pulled down by NAA80 through actin or profilin does exist.

The NAA80 P2 polyproline stretch is a possible reason for the profilin isoform selectivity. PFN1 and PFN2a/PFN2b exhibit a sequence similarity of 61.4%/62.1%, and the differences are reflected in an altered overall positive net-charge for PFN1 and negative charges for PFN2a and PFN2b^19^ While the actin-binding sites of the three isoforms are highly similar, sequence differences mainly occur in the polyproline binding site containing the N- and C-terminal profilin helices, resulting in different affinities towards various polyproline-rich ligands^21,45^. Kursula and colleagues studied the binding of mouse PFN2a to polyproline peptides^20^. While they did not test the sequence of NAA80 polyprolines specifically, they did investigate the interaction between PFN2a and a polyproline-rich ligand from the Formin Homology 1 (FH1) domain of mDia1 using X-ray crystallography. This peptide contains the central binding motif IPPPPPL, which is highly similar to the NAA80 polyproline stretch P2 with the core sequence LPPPPPL. The crystal structure of the PFN2a-mDia1 ligand complex (PDB: 2V8F) revealed that the non-proline residues I and L of the mDia1 peptide serve as anchor to PFN2a, packing against Y6 and W31 of PFN2a. In addition, W3 and F139 were identified as critical residues interacting with the proline residues of the mDia1 peptide^20^. The similarity of the polyproline stretch of mDia1 and NAA80 may indicate a similar interaction interface with PFN2a. Another study using a similar polyproline-rich peptide from human palladin (FPLPPPPPPLPS), demonstrated that both PFN1 and PFN2a were able to bind with similar affinity^46^, which again stands in contrast to the differences observed in our studies. Due to the conservation and presence of the critical residues W31, Y6, and W3 in all three profilin isoforms^19^, the affinity differences for PFN1/PFN2 observed in our experiments is surprising. However, a possible explanation is the aromatic extension of the polyproline binding site by Y29, which is present in PFN2a and PFN2b^19^ This aromatic residue is conserved in PFN2a and PFN2b, but is replaced by S29 in PFN1. Y29 can accommodate an additional proline residue, which may lead to a favored arrangement of the P2 stretch (KGPPLPPPPPLPEC) when bound to PFN2, possibly explaining the specificity of PFN2 interaction with NAA80. In addition, PFN2 exhibits an extensive hydrophobic area in the proximity of Y29, caused by the aliphatic residues I45 and M49^19^ In contrast to the smaller residues A45 and V49 in PFN1, those large residues are protruding from the PFN2 surface and may provide additional anchor points for the NAA80 β6β7 loop.

Our recent study showed that actin-PFN1 is a better substrate for NAA80 than actin alone^33^. Any mechanistic explanation of how PFN2 potentiates NAA80 has to take into account that the potentiation does not just occur when the substrate is full-length actin, but also for actin peptides, which lack the NAA80-interacting residues^33^ and the PFN2 interaction domain present in full-length actin. This suggests a conformational change in NAA80 upon PFN2 binding, rather than just a chaperone effect of PFN2 towards actin. The flexible β6-β7 loop is common to all NATs, forming the peptide substrate-binding pocket along with the α1-α2 loop and residues of the α2 helix. The conformation of the β6-β7 loop is a major determinant of how open the substrate channel is^47^. The extended β6-β7 loop is able to exhibit a multitude of conformations, some of which may hinder substrate or Ac-CoA binding (Fig. 6d). A constraint of the β6-β7 loop upon PFN2 binding may increase substrate accessibility and thereby increase the acetylation rate of NAA80 (Fig. 3c). Alternatively, PFN2 binding to the polyproline stretch P2 might induce conformational changes, possibly propagated via the additional rigid polyproline stretches P1 and P3, and thereby change the positioning of critical residues in the catalytic site or substrate/Ac-CoA binding cleft.

Based on the IP/MS data (Fig. 1 and 4) and the gel filtration complex analysis (Fig. 8), we do not believe that NAA80-PFN2-actin complexes are abundant in human cells. Given the actin:NAA80 ratio of over 3000:1 and the high stoichiometry of actin Nt-acetylation, NAA80 would not be expected to associate tightly with its substrate. Since almost all actin is Nt-acetylated at steady state, we propose that NAA80-PFN2 heterodimers act rapidly on the newly synthesized pool of actin. NAA80-PFN2 dimers are relatively abundant compared to NAA80 monomers (Fig. 8). Thus, preformed NAA80-PFN2 complexes may efficiently associate with monomeric actin and catalyze actin Nt-acetylation. At the same time, it cannot be excluded that actin-PFN2 or actin-PFN1 complexes also may be targeted by NAA80-mediated Nt-acetylation. Indeed, an earlier experiment with actin-PFN1 complexes showed that this heterodimer is more efficiently acetylated by NAA80 than actin alone^33^. We have not performed the corresponding experiment with actin-PFN2, but we note the following four points: 1) PFN2 increases NAA80 activity also towards peptides, demonstrating that interaction between the fold of actin and NAA80 is not required for potentiation. 2) We observe no potentiation from NAA80-PFN1 preincubation towards neither actin peptides nor full-length actin, demonstrating that the effect is PFN2-specific. 3) The lack of evidence of NAA80-actin dimers (Fig. 1b, 2d, 4b-c, 7), indicates that actin-NAA80 affinity *in vivo* is low. 4) NAA80-PFN2 dimers are more prevalent than NAA80-actin-PFN2 heterotrimers (Fig. 8), suggesting that NAA80-PFN2 interaction is neither driven by actin nor dependent on it. Our data support a model of low-affinity, concentration driven PFN2-NAA80 association, where PFN2 recruits newly synthesized actin to NAA80, concurrent with or sequential to ATP binding by actin. We have outlined three possible chains of events in Fig. 9, where newly synthesized actin is processed by NatB (co-translationally) and an unidentified acetylmethionine aminopeptidase (AcMetAP) (either co- or posttranslationally) before recognition by NAA80-PFN2.

**Figure 9:**
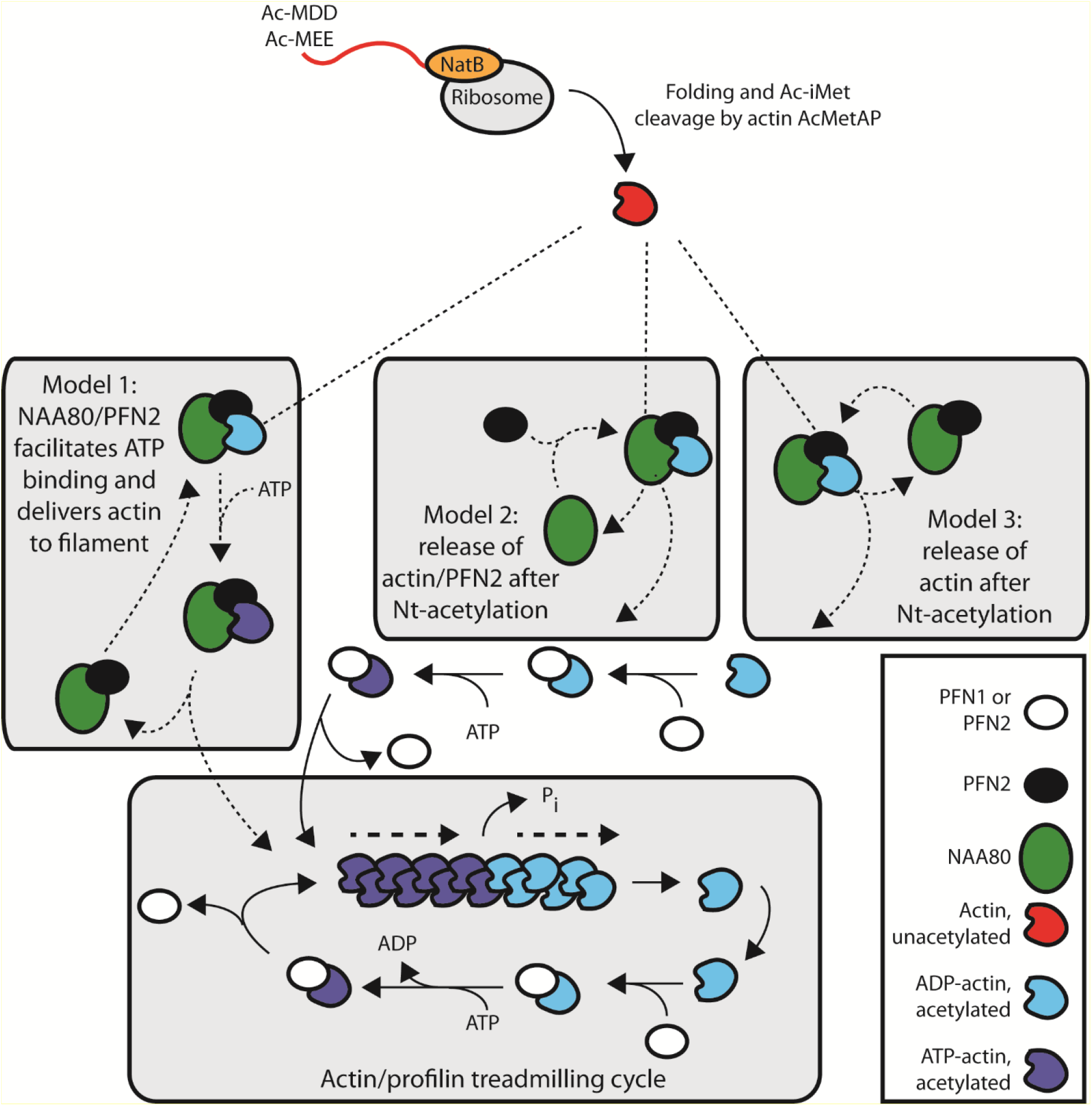
The function of NAA80 and PFN2 in actin N-terminal processing. During synthesis, NatB co-translationally acetylates the actin N-terminus and the acetylated initiator methionine (Ac-iMet) is subsequently removed by acetyl-methionine aminopeptidase (AcMetAP). The NAA80-PFN2 complex Nt-acetylates the actin neo-N-terminus and the Nt-acetylated actin subsequently enters the polymerization pool. Profilins PFN1/PFN2 bind G-actin with high affinity and promote binding of ATP and subsequent polymerization. ADP-actin dissociating at the pointed end binds profilin to reenter the cycle. Three possible options for how the newly synthesized actin enters the polymerization pool are presented: 1) After folding, actin is Nt-acetylated by NAA80-PFN2. NAA80-PFN2 facilitates ATP binding and delivers actin to the filament for polymerization, before being recycled for a new acetylation cycle. 2) After acetylation, NAA80 is released, PFN2 catalyzes actin-ATP binding and actin is delivered to the filament. NAA80 binds PFN2 before a new acetylation cycle. 3) NAA80-PFN2 releases actin after acetylation and is immediately available for a new acetylation cycle. Actin is bound by profilin before ATP binding and delivery to the actin filament.

In conclusion, we have described the direct interaction between the actin NAT NAA80 and PFN2, an important cytoskeletal regulator and actin-binding protein. We hypothesize a constitutive role for NAA80-PFN2 in catalyzing rapid actin Nt-acetylation, which may be adaptive at times of high actin production, for example during cell growth or cell division. The specific roles of different profilins have remained elusive, but we here define a unique role for PFN2 as an enhancer for NAA80-mediated actin acetylation.

## Materials and methods

### Plasmid construction

pcDNA4-V5-NAA80-M23L was constructed by subcloning *NAA80* from pcDNA3.1-NAA80-V5^25^ into the TOPO TA vector pcDNA 4/Xpress-His (Invitrogen). Then the M23L mutation was introduced and the N-terminal Xpress tag was replaced with a V5 tag. Mutations in the vector and *NAA80* were introduced using the Q5 Site-Directed Mutagenesis Kit (New England Biolabs) according to the manufacturer’s protocol. For recombinant expression and subsequent purification PFN1, PFN2a and PFN2b were cloned between the *SapI* and *Eco*RI sites of vector pTYB11 (NEB). NAA80 and its mutants were cloned between the *NdeI* and *Eco*RI sites of vector pTYB12 (NEB). The cDNA of human gelsolin (UniProt ID: P06396) was purchased from Open Biosystems (GE Healthcare). The fragment encoding subdomains 4-6 (residues 434-782) was amplified by PCR and cloned between the *Xho*I and *Eco*RI sites of vector pCold (Takara Bio). The plasmids encoding the profilin isoforms, the gelsolin fragment G4-G6, and the NAA80 wildtype were gifts from Roberto Dominguez (University of Pennsylvania). All primers and plasmids used in this study are listed in Table S6.

### Recombinant protein expression and purification

The pTYB vectors contains a chitin-binding domain, used for affinity purification, and an intein domain, used for self-cleavage and release of the target protein. The proteins cloned in pTYB vectors were expressed in BL21(DE3) cells (Invitrogen), grown in Terrific Broth medium at 37 °C until the OD600 reached a value of 1.5-2, followed by 16 h at 20°C in the presence of 0.4 mM isopropyl-β-d-thiogalactoside (IPTG). Cells were harvested by centrifugation, resuspended in 20mM HEPES pH 7.5, 500mM NaCl, 1mM EDTA, 1 mM 100 μM phenylmethylsulphonyl fluoride (PMSF) (profilin isoforms) or 50 mM Tris pH 8.5, 300 mM NaCl, 1 mM EDTA, 1 mM DTT, 100 μM PMSF (NAA80 constructs) and lysed using a French Press. After clearing the lysate by centrifugation (40,000 x g, 25 minutes, 4°C), the extract was applied onto a self-packed chitin affinity column, and purified according to the manufacturer’s protocol (NEB), followed by additional purification through a HiLoad 16/600 Superdex 75 gel filtration column (GE Healthcare) in 20 mM HEPES pH 7.5, 100 mM NaCl, 1 mM DTT, 1 μM PMSF (profilin isoforms) or 50 mM Tris pH 8.5, 300 mM NaCl, 1 mM EDTA, 1 mM DTT, 100 μM PMSF (NAA80 constructs). Gelsolin from pCold-G4-G6 vector was expressed in Rosetta (DE3) cells, grown in Terrific Broth medium at 37°C until the OD600 reached a value of 0.8 followed by 24 hours at 15°C in the presence of 0.4 mM IPTG. Cells were then harvested by centrifugation, resuspended in 50 mM Tris-HCl pH 8.0, 500 mM NaCl, 5 mM imidazole and 100 μM PMSF, and lysed using a French Press. Gelsolin G4-G6 was purified on a Ni-NTA affinity column.

### Human cell culture and transfection

Wild type HAP1 cells (Horizon C631), *NAA80-KO* (HZGH0003171c012), *PFN1-KO* (HZGHC005831c004), and *PFN2*-KO (HZGHC005248c009) cells were purchased from Horizon Discovery and cultured in Iscove’s Modified Dulbecco’s Medium (IMDM) (Gibco) supplemented with 10% FBS and 1% penicillin-streptomycin at 37 °C and 5% CO2. Prior to experiments, all HAP1 cell lines were passaged until diploid status was confirmed by an Accuri BD C6 flow cytometer using propidium iodide staining. HeLa cells (ATTC CCL-2) were cultured in DMEM supplemented with 10% FBS, 1% penicillin-streptomycin and 2.5 mM *L*-glutamine. Cells were transfected with XtremeGene 9 (Roche) or Fugene (Promega). Cells were harvested 12-48 hours post*-*transfection.

### Purification of endogenous actin from HAP1 cells

Ac-actin (from wildtype HAP1 cells) and non-Ac-actin (from HAP1 *NAA80*-KO cells) were purified using a gelsolin-affinity purification protocol^35^. *NAA80* knockout and wild type HAP1 cells were grown in 15 cm-diameter plates to 70% confluence in IMDM with the addition of 10% fetal bovine serum and 1% penicillin/streptomycin. Cells were harvested and lysed using a glass homogenizer in “Binding buffer” (10 mM Tris-HCl pH 8.0, 5 mM CaCl_2_, 1 mM ATP, 1 mM TCEP) supplemented with cOmplete protease inhibitor cocktail (Roche), 25 nM calyculin A, 1 x HALT protease inhibitor cocktail (ThermoFisher Scientific), and 1 mM PMSF. Cell lysates were incubated overnight at 4 °C by gentle mixing with an affinity tag consisting of gelsolin subdomains G4 to G6, which binds actin in a Ca^2+^-dependent manner. The lysates were centrifuged for 45 minutes at 100,000 x *g* and the supernatant was incubated for 1 hour with 5 mL Ni-NTA resin containing HisTrap FF column (GE Healthcare), which binds the gelsolin-actin complex through an N-terminal polyhistidine affinity tag on the gelsolin fragment. The resin was washed with ten column volumes of Binding buffer with the addition of 100 mM KCl and 20 mM imidazole, followed by five column volumes of Binding buffer without CaCl_2_. Actin was then eluted with 4 mL of Binding buffer in which CaCl_2_ was replaced by 1 mM EGTA. The released actin was polymerized for 1 hour at 25 °C with the addition of 1 mM MgCl2 and 100 mM KCl. The polymerized actin was then pelleted by centrifugation for 1 hour at 270,000 x *g*, sheared using a glass homogenizer to break the pellet, and depolymerized through a three-day dialysis against G-actin buffer (2 mM Tris-HCl pH 8.0, 0.2 mM CaCl_2_, 0.2 mM ATP), followed by centrifugation for 1 hour at 270,000 x *g* to remove any actin that did not depolymerize.

### DTNB N-terminal acetylation assay

The 5,5’-dithiobis-2-nitrobenzoic acid (DTNB) Nt-acetylation assay was performed as described^34^ Purified enzymes (300 nM) were mixed with synthetic peptides (300 μM) and Ac-CoA (300 μM) in acetylation buffer (50 mM Tris-HCl pH 8.5, 200 mM NaCl, and 2 mM EDTA) at 37 °C, and reactions were quenched after 60 minutes with quenching buffer (3.2 M guanidinium-HCl, 100 mM Na_2_HPO_4_, pH 6.8). To measure CoA production, DTNB (2 mM final, dissolved in 100 mM Na2HPO4, pH 6.8 and 10 mM EDTA) was added to the quenched reactions. The thiol present in the enzymatic product, CoA, cleaves DTNB and produces 2-nitro-5-thiobenzoate (TNB^2-^), which is readily quantified by monitoring the absorbance at 412 nm. Background absorbance was determined in negative controls (enzyme added after quenching buffer) and subtracted from the absorbance determined in each individual reaction. Thiophenolate production was quantified assuming ε = 13.7 × 103 M^-1^ cm^-1^. For the profilin activation assays with full-length NAA80, the indicated molar ratio of profilin (either PFN1, PFN2a or PFN2b) or of bovine serum albumin (BSA) was added to the reaction mix. Product formation was normalized to the product formation in the corresponding BSA reaction. The normalized standard deviation was calculated using the formula 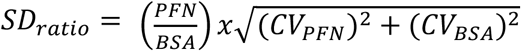, where 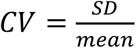 of the PFN or BSA normalized product formation (n=4 for each molar ratio). For the activation assays with NAA80 mutants (ΔP123 and polyGS2), the NAA80/profilin activity is normalized to the activity of NAA80 with no added profilin. Ratio standard deviation was calculated in a similar manner as above: 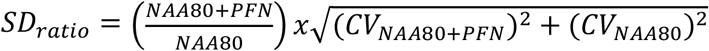 where 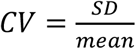 of the NAA80/profilin or NAA80 alone normalized product formation (n = 3 for each molar ratio).

### Analytical ultracentrifugation (AUC)

Sedimentation velocity analysis of ATTO-488 (ATTO-TEC) labeled profilins was performed with a ProteomLab Beckman XL-A analytical ultracentrifuge (Beckman Coulter, Brea, California) equipped with an AVIV fluorescence detection system (Aviv Inc., Lakewood, USA). Unlabeled components were added at various concentrations, as indicated in the plots, to 500 nM profilin. NAA80-profilin complexes were measured in 20 mM HEPES, 100 mM NaCl, 1 mM EDTA, 1 mM DTT, pH 8.5. For measurements including actin (human platelet actin, 99% pure, from Cytoskeleton, Inc., catalog nr: APHL99), the buffer was 5 mM Tris, 0.2 mM CaCl_2_, 0.2 mM ATP, 5% sucrose, pH 8.0. Ultracentrifugation was carried out with 42,000 rpm at 20 °C and dF/dt plots were generated by subtracting scans from a selected time range and normalizing the data against the initial fluorescence intensity. All plots were generated from samples measured in the same experiment to ensure identical sample handling.

### Peptide synthesis and quality validation

The peptides were synthesized by means of the simultaneous multiple peptide synthesis (Schnorrenberg 1989 Tetrahedron) on the following instrument: SYRRO (MultiSynTech, Germany), using the Fmoc/But strategy by -SHEPPARD (Fmoc Solid State Synthesis 1999 Book). Couplings were performed using 3-6 equivalents Fmoc-amino acid/TBTU and 6-12 equivalents N-methylmorpholine on the following resin: Tentagel S Trityl resin (RAPP Polymere, Germany), loading: 0.20 mmol/g resin. Peptides were synthesized as C-terminal acids (-COOH). The following protection groups have been used: Cys(trt), Arg(Pbf), Ser(But), Thr(But), Asp(OBut), Glu(OBut), Asn(Trt), Gln(Trt), Lys(Boc), His(Trt), Trp(Boc). Peptides were deprotected and cleaved from resin by trifluoroacetic acid (TFA)/thioanisole/thiocresol (95:2.5:2.5), adding 3% triisopropylsilane for 3 hours at RT. To purify the synthesized peptides following preparative HPLC has been used: HPLC instrument Shimadzu LC-8A with UV/VIS detector SPD-10A. Column: Ultrasep ES (RP-18), 10 μm, 250 x 20 mm. Solvent: A = 0.05% TFA in water, solvent B = 0.05% TFA in 80% ACN/water with a gradient of 30 minutes, flow rate: 15 ml/minute, detection: 220nm. For validation purposes the following analytical HPLC has been used: HPLC instrument Shimadzu LC-30AD with photodiode array detector SPD-M20A; column Ascentis Express Peptide ES-C18, 2.7 μm, 30 x 2.1 mm (Supelco, USA).

Solvent was used as described above with a flow rate of 0.5 ml/minute and a linear gradient of 21.4% B/minute; detection at 220 nm. Peptides were characterized by MALDI-TOF by means of a MALDI Axima Assurance instrument (Shimadzu, Japan) in linear mode. Detected molecular weight as [M+H]^+^ or M+22 [M+Na]^+^. The peptides were lyophilized in form of the TFA salt and resuspended in water to a stock solution of 5 mM.

### Immunoprecipitation (IP) experiments

HAP1 cells (wild-type (WT), *PFN1-KO* or *PFN2*-KO) were split 1:3 and the next day transfected using XtremeGene 9 (Roche) with 5 μg V5-NAA80 (M23L) or lacZ-V5 plasmid for 48 hours. Cells were harvested by trypsinization and washed twice in ice-cold PBS. They were lysed by resuspending cell pellets in 12 μl IPH buffer (50 mM Tris-HCl, pH 8.0, 150 mM NaCl, 1 % Nonidet-P 40) supplied with 1 tablet/50 mL c0mplete EDTA-free Protease Inhibitor Cocktail (Roche), per mg cell pellet and rotating on a wheel at 4°C for 15 minutes. Lysates were cleared by centrifugation at 17,000 x *g* for 5 minutes and used for immunoprecipitation. 2 μg anti-V5 was added to the lysates and they were incubated with rotation at 4°C for 2 hours, before 15 μl Dynabeads Protein G magnetic beads (Invitrogen) were added. The bead complexes were left to form overnight at rotation and 4°C. The immune complexes were retrieved by removing the supernatant on a magnet and washing 3 times with 500 μl IPH buffer. The beads were resuspended in 1x sample buffer (Bio-Rad), boiled, and the supernatant was loaded on a gel and probed with the indicated antibodies.

### Circular dichroism spectroscopy

Circular dichroism spectroscopy (CD) experiments were conducted using a Jasco J-810 spectropolarimeter. Data was collected from 0.1 mg/ml protein samples in 10 mM Na-phosphate buffer pH 8.0, 100 mM NaF, 1 mM TCEP. Wavelength scans (185 nm – 280 nm, 1 nm data pitch, 3 accumulations) were performed at 20 °C. Baselines were subtracted from sample spectra and raw data were smoothened in GraphPad Prism 8 using a polynomial function of 2^nd^ order with a smoothing window of 5. Thermal denaturation CD data were collected from 0.2 mg/ml protein in 1 mM TCEP (20 °C – 95 °C, 0.2 °C data pitch) in triplicate. The data were fitted into a sigmoidal function in GraphPad Prism 8 and the melting temperature *(T_m_)* was extracted from the midpoint of the curve.

### Differential scanning fluorimetry

Protein thermal stability was assessed by differential scanning fluorimetry (DSF). Measurements were conducted using 0.1 mg/ml protein in 10 mM HEPES pH 8.0, 100 mM NaCl, 5x SYPRO Orange (Invitrogen). Fluorescence was followed during protein denaturation from 20 °C to 98 °C, with a data pitch of 2 °C/minute. All measurements were performed in triplicate. Data were fitted into a sigmoidal function in GraphPad Prism 8 and the *T_m_* was extracted from the midpoint of the curve.

### Multi-angle light scattering

Size exclusion chromatography-coupled multi-angle light scattering (SEC-MALS) was used to analyze protein monodispersities and molecular weights. SEC was performed using an Äkta Purifier (GE Healthcare) and a Superdex 75 Increase 10/300 GL column (GE Healthcare) in 20 mM HEPES pH 8.5, 100 mM NaCl, 2 mM EDTA. For each measurement, 200 – 300 μg of protein was injected and gel filtrated at a flow rate of 0.5 ml/minute. Light scattering was recorded using a miniDAWN TREOS instrument (Wyatt Technology). Protein concentration in each elution peak was determined using differential refractive index (dRI). The data were analyzed using the ASTRA 6.2 software (Wyatt Technology).

### Small-angle X-ray scattering

SAXS measurements were performed at the EMBL P12 beamline at PETRA III, DESY (Hamburg, Germany) and at the B21 beamline at Diamond Light Source (Oxfordshire, UK). Measurements of NAA80 variants and ternary complexes were conducted in SEC-SAXS mode, using a Superdex 200 Increase 10/300 GL column (GE Healthcare). Gel filtration was performed at a flow rate of 0.5 ml/minute (20 mM HEPES pH 8.0, 100 mM NaCl, 1 mM EDTA) at 10 °C. Single proteins (NAA80 and NAA80-ΔP123) were injected at a concentration of 5.8 mg/ml and 10 mg/ml, respectively. Individual components of the protein complex, NAA80, PFN2a/b, human platelet actin (Cytoskeleton Inc., APHL99), were mixed in equimolar ratios at 50 μM and incubated on ice for 30 minutes prior to measurement.

SAXS measurements from 1.1 – 5.7 mg/ml PFN1, PFN2a, and PFN2b samples were performed in batch mode at 10 °C. The sample buffer contained 20 mM HEPES pH 8.0, 100 mM NaCl, and 1 mM DTT. Data reduction, processing, and analysis were performed using the ATSAS 2.8 package^48^. SEC-SAXS frames were analyzed using CHROMIXS^49^ *Ab initio* models were generated using GASBOR^50^. Missing loops and termini were built in rigid bodies using CORAL^51^ and ensemble optimization analysis was performed using EOM^52,53^. CRYSOL was used to calculate theoretical scattering profiles based on crystal structures^54^ Data processing, analysis, and modelling details are listed in Table S4 and S5.

### Protein disorder prediction

The NAA80 sequence (UniProt ID: Q93015) was submitted to the IUPred2A server^55^, predicting disordered protein regions (IUPred2) and disordered binding sites (ANCHOR). The output graph represents the disorder tendency of each individual residue, with higher scores corresponding to higher disorder probabilities.

### Actin reacetylation assay in NAA80-KO cells

Degree of actin acetylation after transfection with a NAA80 variant was measured essentially as described^33^. Wildtype NAA80 rapidly reacetylates β-and γ-actin, necessitating a short transfection time to discriminate between the efficiency of different NAA80 variants. We have analyzed cells 8-14 hours post-transfection, since after 24 hours most variants would have completely Nt-acetylated the cellular actin pool. 7 x 10^6^ HAP1 *NAA80-KO* cells (50% confluent) were transfected with 3 μg construct harboring the human full-length NAA80 (V5-HsNAA80 (M23L)), and 1.8 μg of NAA80-ΔP123 (V5-HsNAA80 (M23L) Δ238-284). Cells were harvested and lysed in IPH buffer as described above. Lysates were analyzed by Western blotting using anti-Ac-β-actin, anti-Ac-γ-actin, anti-pan-actin, and anti-V5. Degree of actin acetylation relative to V5 expression was calculated from 4 independent experiments by dividing the Ac-β- or Ac-γ-actin signals by the corresponding V5 signals. Standard deviations for the ratios were calculated using the following formula: signals (n = 4). Significance was calculated using a two-way ANOVA, implemented in Graphpad Prism v. 8.4.2., with significance threshold 0.05.

### Immunoprecipitation for MS

Immunoprecipitation was performed essentially as described above. 10 cm dishes of HeLa or HAP1 cells were transfected in triplicates with 5 μg NAA80-V5, V5-NAA80 (M23L), V5-NAA80 (M23L) Δ238-284, or lacZ-V5 for 48 hours. Cells were harvested by trypsinizing, lysed in IPH buffer and lysates were cleared for 5 minutes at 17,000 x *g*. Immunoprecipitation with PureProteome A/G magnetic beads (Merck Millipore) and anti-V5 (Invitrogen) proceeded overnight. Bound proteins were eluted by boiling in 50 μL FASP lysis buffer (2% SDS, 100 mM Tris-HCl, pH 7.6, 100 mM DTT) for 5 minutes. Beads were separated on the magnet and the eluate was collected and used for FASP, as described below.

### Filter-aided sample preparation (FASP) and stagetips desalting

Filter-aided sample preparation (FASP) was performed on the V5 immunoprecipitates essentially as described^56^. Eluted proteins were mixed with UA (8 M urea, 100 mM Tris-HCl, pH 8.0) transferred to Microcon 30 kDa centrifugal filter units and cysteine-alkylated with 50 mM iodoacetamide. The buffer was exchanged to 50 mM ammonium bicarbonate by sequential centrifugation, proteins were trypsinized overnight (Sequencing Grade Modified Trypsin, Promega, cleaves C-terminal to lysine and arginine residues, except when followed by a proline) and the tryptic peptides were recovered by centrifugation. The peptides were dried in a vacuum evaporator, resuspended in 1% formic acid, and desalted using in-house packed stagetips essentially as described^57^ At each stage of the protocol, the stagetips were spun at around 1500 x *g* for 1-2 minutes, until almost no solution remained. They were first activated with 200 μl acetonitrile with 1% formic acid, then equilibrated three times with 200 μl 1% formic acid. The peptides were loaded and the flowthrough was run through the stagetip a second time. The stagetip was then washed three times with 1% formic acid, before the peptides were eluted with rounds of 200 μl 80% acetonitrile in 1% formic acid. The eluate was vacuum dried and resuspended in A* buffer (5% acetonitrile and 0.1% trifluoroacetic acid), and the peptide concentration was measured by absorbance at 280 nm using a Nanodrop spectrophotometer (Thermo Fisher).

### Liquid chromatography-mass spectrometry

0.5 μg peptides in A* buffer were injected into an Ultimate 3000 RSLC system (Thermo Scientific) connected to a Q-Exactive HF mass spectrometer (Thermo Scientific) with EASY-spray nano-electrospray ion source (Thermo Scientific). The sample was loaded and desalted on a pre-column (Acclaim PepMap 100, 2 cm x 75 μm ID nanoViper column, packed with 3 μm C18 beads) at a flow rate of 5 μl/minute for 5 minutes with 0.1% TFA. Peptides were separated in a biphasic acetonitrile gradient from two nanoflow UPLC pumps (flow rate of 200 nl/minute) on a 50 cm analytical column (PepMap RSLC, 50 cm x 75 μm ID EASY-spray column, packed with 2 μm C18 beads). Solvent A and B were 0.1% TFA (vol/vol) in water and 100% acetonitrile respectively. The gradient composition was 5% B during trapping (5 minutes) followed by 5-8% B over 0.5 minutes, 8-24% B for the next 109.5 minutes, 24-35% B over 25 minutes, and 35-90% B over 15 minutes. Elution of hydrophobic peptides and column conditioning were performed by isocratic elution for 15 minutes with 90% B and 20 minutes of isocratic conditioning with 5% B. The total LC run time was 195 minutes. The eluting peptides were ionized in the electrospray and analyzed by the Q-Exactive HF. The mass spectrometer was operated in the DDA-mode (data-dependent-acquisition) to automatically switch between full scan MS and MS/MS acquisition. Instrument control was through Q Exactive HF Tune 2.4 and Xcalibur 3.0. Survey full scan MS spectra (from m/z 375-1500) were acquired in the Orbitrap with resolution R = 120,000 at m/z 200, automatic gain control (AGC) target of 3 x 10^6^ and a maximum injection time (IT) of 100 ms. The 15 most intense eluting peptides above an intensity threshold of 50,000 counts, and charge states 2 to 6, were sequentially isolated to a target value (AGC) of 105 and a maximum IT of 100 ms in the C-trap, and isolation with maintained at 1.6 m/z (offset of 0.3 m/z), before fragmentation in the HCD (Higher-Energy Collision Dissociation) cell. Fragmentation was performed with a normalized collision energy (NCE) of 28%, and fragments were detected in the Orbitrap at a resolution of 15,000 with first mass fixed at m/z 100. One MS/MS spectrum of a precursor mass was allowed before dynamic exclusion for 20 seconds with “exclude isotopes” on. Lock-mass internal calibration (m/z 445.12003) was enabled. The spray and ion-source parameters were as follows: ion spray voltage = 1800 V, no sheath and auxiliary gas flow, and a capillary temperature of 260 °C.

### Database searching, filtering and statistics for MS experiments

The resulting RAW files were processed using MaxQuant (version 1.6.1.0) and the integrated search engine Andromeda^58,59^. They were searched against a human proteome database containing 20,352 annotated, canonical and isoform entries, retrieved from UniProt on 6 Jan 2020, and a reverse decoy database automatically generated by the search engine. Carbamidomethylation of cysteine was set as a fixed modification, and N-terminal protein acetylation and methionine oxidation as variable modifications. Protease specificity was set as trypsin (C-terminal to lysine and arginine, except when followed by a proline). Minimum peptide length was set to 7 amino acids and maximum peptide mass was set to 4600 Da. Peptide and protein identifications were filtered to a 1% false discovery rate (FDR). The MaxLFQ label-free quantification algorithm^60^ was used to obtain relative quantities of each protein group in the different samples. LFQ settings were as follows: minimum ratio count: 2; “Fast LFQ” was checked; and “Skip normalization” was not checked. Resulting protein group lists were processed in Perseus (version 1.6.10.43)^61^. Proteins identified only by reverse sequences and by modified sites were removed. Proteins enriched by NAA80 IP were identified by making a volcano plot, with a S0 of 0.1 and FDR of 0.05. The mass spectrometry proteomics data have been deposited to the ProteomeXchange Consortium via the PRIDE^62^ partner repository (https://www.ebi.ac.uk/pride/archive/) with the dataset identifier PXD020188.

### Gel filtration complex assay

4 x 10 cm dishes of HAP1 cells were harvested and lysed in IPH buffer as described above. Cell lysate was ultracentrifuged for 20 minutes at 100,000 x *g* to clear the lysate from F-actin and other insoluble structures. The supernatant was subjected to gel filtration using a preequilibrated Superdex 200 Increase 10/300 GL column at a flow rate of 0.5 ml/minute. The gel filtration buffer was composed of 50 mM Tris-HCl (pH 8.0), 200 mM NaCl and 10% glycerol. 0.5 ml fractions were collected, precipitated with TCA/chloroform, and the pellets were dissolved by boiling in 1 x SDS-PAGE loading buffer. The proteins from each fraction were separated by SDS-PAGE and analyzed by Western blotting. The calibration mix (Protein Standard Mix 15 – 600 kDa, Merck Millipore) was composed of the following proteins: 0.5 g/l bovine thyroglobulin (MW ~ 670 000), 1.0 g/l γ-globulins from bovine blood (MW ~ 150 000), 1.0 g/l grade VI chicken egg albumin (MW ~ 44 300), 1.0 g/l Ribonuclease A type I-A from bovine pancreas (MW ~ 13 700) and 0.01 g/l *p*-aminobenzoic acid (pABA).

## Acknowledgements

We thank Henriette Aksnes and Nina Glomnes for experimental inputs and technical assistance. The gelsolin, NAA80 and profilin plasmids were kind gifts from Prof. Roberto Dominguez (University of Pennsylvania). SAXS measurements were performed in collaboration with the research group of Prof. Petri Kursula (University of Bergen, Norway). We gratefully acknowledge Dr. Arne Raasakka and Prof. Petri Kursula for guidance with SAXS data collection, analysis, interpretation, and presentation. We acknowledge Dr. Juha Kallio for assistance with SEC-MALS experiments. CD, DSF, and SEC-MALS experiments were performed at the Biophysics, Structural Biology and Screening (BiSS) facility at the University of Bergen. LC/MS runs were performed at PROBE, the proteomics core facility at the University of Bergen. We are grateful for supportive synchrotron beamline staff at PETRAIII EMBL P12 and Diamond Light Source B21 beamlines. This work has been supported by iNEXT, grant number 653706, funded by the Horizon 2020 programme of the European Commission. The Arnesen lab was supported by funds from the Research Council of Norway (Project 249843), the Norwegian Health Authorities of Western Norway (Project 912176), the Norwegian Cancer Society, and the European Research Council (ERC) under the European Union Horizon 2020 Research and Innovation Program under grant agreement 772039.

