## Supplemental figures and Supplemental tables 3-6 for "PFN2 and NAA80 cooperate to efficiently acetylate the N-terminus of actin"

```
PFN2a (117-126)  --------EGVHGGGLNK--------------
PFN2a (109-116)  VLVFVMGK------------------------
PFN2a (109-140)  VLVFVMGKEGVHGGGLNKKAYSMAKYLRDSGF
PFN2b (109-140)  ALVIVMGKEGVHGGTLNKKAYELALYLRRSDV
PFN2b (109-115)  ALVIVMGK------------------------
PFN2b (117-127)  --------EGVHGGTLNKK-------------
PFN2b (127-136)  ------------------KAYELALYLR----
PFN2b (128-136)  -------------------AYELALYLR----
```

**Supplemental figure 1:** PFN2 isoform specific peptides identified in at least one NAA80-V5 IP aligned with the variable region of PFN2a and PFN2b (aa 109-140). The differing sites are labeled with blue (PFN2a) or red (PFN2b).

```
Homo_sapiens_|Q93015|                     1 MQELTLSPGPAKLTPTLDPTHRMELILSTSPAELTLDPACQPK-----------------  43
Mus_musculus_|Q9R123|                     1 ----------------------MELILSTSPAKLTLDPARQPELTLRFNLSKLTLDPARQ  38
Rattus_norvegicus_|A0A0G2JV35|            1 ----------------------MELILSTSPAKLTLDPACQPELTLRFNLTKLTLDPARQ  38
Danio_rerio_|E7FBQ5|                        ------------------------------------------------------------
Takifugu_bimaculatus_|A0A4Z2B4F3|         1 -------------------------------------------------MSETLTS-S--   8
Oryzias_latipes_|A0A3P9HTG7|              1 -------------------------------------------------MSEATGPPN--   9
Portunus_trituberculatus|A0A5B7DRL4|        ------------------------------------------------------------
Armadillidium_vulgare_|A0A444SNE1|          ------------------------------------------------------------
Drosophila_melanogaster_|Q59DX8|            ------------------------------------------------------------
Anopheles_gambiae_|F5HLV5|                  ------------------------------------------------------------
Caenorhabditis_elegans_|Q09518|             ------------------------------------------------------------

Homo_sapiens_|Q93015|                    44 -----------LPLDSTCQPEMTFNPGPTELTLDPEHQPEETPAPSLAELTLEPVHRRPE  92
Mus_musculus_|Q9R123|                    39 PELSLSPRLAELTLDPTCHPEMSLSPGPAELTLDPQHQAKELPVPKLPELILEPVHCRPE  98
Rattus_norvegicus_|A0A0G2JV35|           39 SELSLSPRLAELTLDSTCHPEMSLSPGPAELTGDPQHQAKESLVPKLAELTLEPVHCRPE  98
Danio_rerio_|E7FBQ5|                      1 -------------------------------------MCD--GDVSCTVFRIEPLHERWD  21
Takifugu_bimaculatus_|A0A4Z2B4F3|         9 ----PANRG---------E-----------VCVHPRTSAS--DCGHPENVHIAPIHLRPD  42
Oryzias_latipes_|A0A3P9HTG7|             10 ----PKHPTNLPFVPDCG------------DHVDGEEPCT--DSEQPPGVRLVPVHQRPD  51
Portunus_trituberculatus|A0A5B7DRL4|      1 -------------------------------------------MKGEEGLRLVTLHSHPE  17
Armadillidium_vulgare_|A0A444SNE1|        1 ---------------------------------------------MKEQLKLLPLHKHPE  15
Drosophila_melanogaster_|Q59DX8|          1 ---------------------MRYIKSEPYYEGLP------PFNVSGSPFNVVPIHNYPE  33
Anopheles_gambiae_|F5HLV5|                1 ------------------------------MTLPP------PQPVISEPYKVVPIHKHRE  24
Caenorhabditis_elegans_|Q09518|           1 ----------------------------------------------MPDLFFVTLYDRQD  14

Homo_sapiens_|Q93015|                    93 LLDACADLINDQWPRSRTSRLHSLGQSSDAFPLCLMLLSPHPTLEAAPVVVGHARLSRVL 152
Mus_musculus_|Q9R123|                    99 LMSACADLINDQWPRSRASRLHSLGQSSDAFPLCLMLLSPQPTPGAAPVVVGHARLSRVL 158
Rattus_norvegicus_|A0A0G2JV35|           99 LMSACADLINDQWPRSRASRLHSLGQSSDAFPLCLMLLSPQPTPGAAPIVVGHARLSRVL 158
Danio_rerio_|E7FBQ5|                     22 LEEACAQLLNDQWQRSMGARIHSLHQSSHDYPVCLLLLQGERQTQ-HEKVIGHARLSRVL  80
Takifugu_bimaculatus_|A0A4Z2B4F3|        43 LLVPCADLVNSEWQRSQAARVHSLMKSCQDFPICLVLLQGPPE---GERLLGHSRLSRVV  99
Oryzias_latipes_|A0A3P9HTG7|             52 LLVPCADLVNSEWQRSQAARVHALQKSCSEFPVCLVLLLGHKG---TERLLGHARLSRVV 108
Portunus_trituberculatus|A0A5B7DRL4|     18 HTEECMKILNDQWPRSRTMRMRSLSTSCDQFPTSLLLLKTTSQ--GDTEVIGHSRLNTLP  75
Armadillidium_vulgare_|A0A444SNE1|       16 FKDACVEMLNKEWPRNVTLRCRFLDSSCDDLPTCLILLLTDEN--GSDILVGHSKLTVIY  73
Drosophila_melanogaster_|Q59DX8|         34 LMKDTCALINAEWPRSETARMRSLEASCDSLPCSLVLTTEGM-----CRVIAHLKLSPIN  88
Anopheles_gambiae_|F5HLV5|               25 LMDQCIALINSEWPRSYTARLWSLESSKETLPTSFVLTTTDIK--DETIVLAHAKLSPIP  82
Caenorhabditis_elegans_|Q09518|          15 LLKESMTFLNSEWPRSDGSREHSQKKSCRQSPPMSFLLLNKE----NDEILGHSRITHLP  70

Homo_sapiens_|Q93015|                   153 NQPQSLLVETVVVARALRGRGFGRRLMEGLEVFAR-ARGFRKLHLTTHDQVHFYTHLGYQ 211
Mus_musculus_|Q9R123|                   159 DQPHSLLVETVVVARPLRGRGFGRRLMEGLEAFAR-ARGFRRLHLTTHDQLYFYAHLGYQ 217
Rattus_norvegicus_|A0A0G2JV35|          159 DHPHSLLVETVVVARALRGRGFGRRLMEGLEAFAR-ARGFQRLHLTTHDQLYFYAHLGYQ 217
Danio_rerio_|E7FBQ5|                     81 GS-RSLLVESVVVCKSLRGKGYGRILMEGVERYAK-GRGCTRLCLTTHDKQHFYAHLGFV 138
Takifugu_bimaculatus_|A0A4Z2B4F3|       100 GQSSSLFVESVVVSKAERGRGYGRTLMEQTERYAR-RRGFRRLCLTTHDKQHFYAHLGYV 158
Oryzias_latipes_|A0A3P9HTG7|            109 GHGGSLFVESVVVSKEERGKGYGRVLMEETERYAR-RRGFKRLCLTTHDKQHFYAHLGYV 167
Portunus_trituberculatus|A0A5B7DRL4|     76 REPDAAWIESVVIRRDLRGAGYGRQLMTRTEEYAR-VSGFTTMYLSTHDQQVFYGKLGYE 134
Armadillidium_vulgare_|A0A444SNE1|       74 NEENSVFIESVIIHHDYRGRGFGKVLMSKTEEYAA-NLGFKTIVLNTK-LKGFYSKLGYK 131
Drosophila_melanogaster_|Q59DX8|         89 SKKKACFVESVVVDKRHRGQGFGKLIMKFAEDYCRVVLDLKTIYLSTIDQDGFYERIGYE 148
Anopheles_gambiae_|F5HLV5|               83 ADREAVFVESVVVTRERRGQGIGRLLMQEVEKHCFHKLHLKKIYLSTIDQQAFYAKLGYK 142
Caenorhabditis_elegans_|Q09518|          71 NRDHALWIESVMIKKDQRGLGLGKFLMKSTEKWMT-EKGFNEAYLSTDDQCRFYESLGYE 129

Homo_sapiens_|Q93015|                   212 LGEPVQGLVFTSRRLPATLLN----AFPTAPSPRP------PRK--APNLTAQAA----- 254
Mus_musculus_|Q9R123|                   218 LGEPVQGLAFTNRRLSTTVLR----AFSKPPCPQP------PCK--EPILAAQAV----- 260
Rattus_norvegicus_|A0A0G2JV35|          218 LGEPVQGLAFTNRRLPTNVLR----AFSKPPCPQP------PCK--APVLAAQAI----- 260
Danio_rerio_|E7FBQ5|                    139 LSKPVQSVGTLASFMPMEILH----RFCRTAENEE------EER--KFKVTNHAK-S--- 182
Takifugu_bimaculatus_|A0A4Z2B4F3|       159 LSTPVQSAGAMTTFVPMETLL----RFSAIPNTGA------RNS--Q---GSRKS-SGGS 202
Oryzias_latipes_|A0A3P9HTG7|            168 LSAPVQNTGPVMALVPMAMLM----RLSRVPDDQK------AAN--TKTEASQES-SGGS 214
Portunus_trituberculatus|A0A5B7DRL4|    135 FCPPVCIYGGSVNKHLI---PKH---FIPPQLTQ----------------GVQALSLKNS 172
Armadillidium_vulgare_|A0A444SNE1|      132 FSEPVCLIRHNASKALA---HTFKYKFNSSEYNKNCTLNSNNCNSLHFDNSKTPLALKLS 188
Drosophila_melanogaster_|Q59DX8|        149 YCAPITMYGPRHCELP-------------------------------------------- 164
Anopheles_gambiae_|F5HLV5|              143 LCSAINIFGSRPTL---------------------------------------------- 156
Caenorhabditis_elegans_|Q09518|         130 KCDPIVHSTTATC--IF---PAM---------N-------------HFQNAAASNPSFLS 162

Homo_sapiens_|Q93015|                   255 -------------------------PRGP------------------KGPPLPPPPPLPE 271
Mus_musculus_|Q9R123|                   261 -------------------------PRSS------------------KGPPLPPPPPLPQ 277
Rattus_norvegicus_|A0A0G2JV35|          261 -------------------------PRSS------------------KGPPLPPPPPLP- 276
Danio_rerio_|E7FBQ5|                    183 -----TPSVLP-----------------PAPPPPPPPPQI-------YSSPPPPQPPI-- 211
Takifugu_bimaculatus_|A0A4Z2B4F3|       203 PPDIPTNSSLPPPPPPPPLPNSQSALLLPPPPPPPPPPPLPNSQSALLPPPPPPPPPLPN 262
Oryzias_latipes_|A0A3P9HTG7|            215 LLA-------------------------THPPPPPS-----------IPPPPPP------ 232
Portunus_trituberculatus|A0A5B7DRL4|    173 VVKDQKQGAPP-----------------------------------------P------- 184
Armadillidium_vulgare_|A0A444SNE1|      189 ST---TDSSTP-----------------------------------------P------- 197
Drosophila_melanogaster_|Q59DX8|            ------------------------------------------------------------
Anopheles_gambiae_|F5HLV5|                  ------------------------------------------------------------
Caenorhabditis_elegans_|Q09518|         163 KIAQPSASSTV-----------------------------------------S------- 174

Homo_sapiens_|Q93015|                   272 CLTISPPVP--------------SGPPSKSLLETQYQNVRGRPIFWMEKDI----      308
Mus_musculus_|Q9R123|                   278 SLTASPPPS--------------PEPLPQSPLETCYRDLKGCPIFWMEKDI----      314
Rattus_norvegicus_|A0A0G2JV35|          277 -LTTSPPPS--------------PEPLPQSPLETRYRDLKGCPIFWMEKDI----      312
Danio_rerio_|E7FBQ5|                    212 --------SCPP-PPPPPPPPLFCAPVSPTLEQTPYTDNSGLPIFWMHKDI----      253
Takifugu_bimaculatus_|A0A4Z2B4F3|       263 SQSALLPPPPPP-PPPPPPPRSTAQPEVQTLTETPYRDARGVPIFWMHKDI----      312
Oryzias_latipes_|A0A3P9HTG7|            233 -SNPHPPPPPPP-PPPPPPPKCGEQLVVQTLTETPYRDAKGNPIFWMHKDI----      281
Portunus_trituberculatus|A0A5B7DRL4|    185 ---ATPAPPHPSLPRPPPLPTS--SK-LTKDHITKMEP-STLTKMYMTKEL----      228
Armadillidium_vulgare_|A0A444SNE1|      198 ---SLPPPPPPP-PPPPSRPVTFRNR-SVLNESKDLYPLEENVHIFMRKAITY--      245
Drosophila_melanogaster_|Q59DX8|        165 -------------------------------------SLQNAKKKYMKKVL----      178
Anopheles_gambiae_|F5HLV5|              157 -----------------------------------NKS---TKKIWMFKSADS--      171
Caenorhabditis_elegans_|Q09518|         175 ---ASAPPPPP---PPPMAPK------MVTRSTSPIVDVNTIDHQYMRKWLKPTE      217
```

**Supplemental figure 2: NAA80 alignment.** NAA80 protein sequences from *Homo sapiens* (Q93015), *Mus musculus* (Q9R123), *Rattus norvegicus* (A0A0G2JV35), *Danio rerio* (E7FBQ5), *Takifugu bimaculatus* (A0A4Z2B4F3), *Oryzias latipes* (A0A3P9HTG7), *Portunus trituberculatus* (A0A5B7DRL4), *Armadillidium vulgare* (A0A444SNE1), *Drosophila melanogaster* (Q59DX8), *Anopheles gambiae* (F5HLV5) and *Caenorhabditis elegans* (Q09518). Sequences were retrieved from UniProt (identifiers in parentheses) and aligned with Clustal Omega.

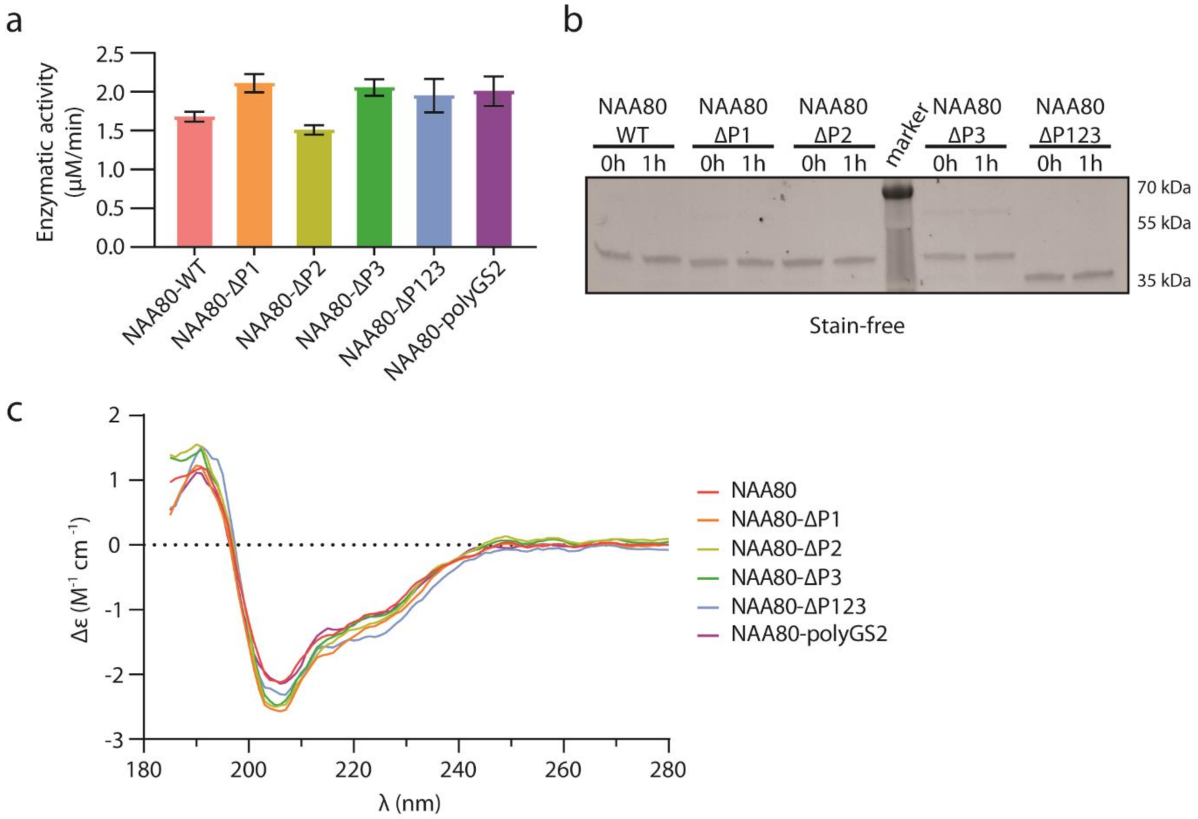

**Supplemental figure 3: Basal activity and stability of NAA80-WT and polyproline mutants.** a) Enzymatic activity of the indicated NAA80 variants were measured using the DTNB assay (n = 4 for all variants). b) Enzymes used in S3a were resolved on an SDS-PAGE gel. c) CD spectra for NAA80 variants.

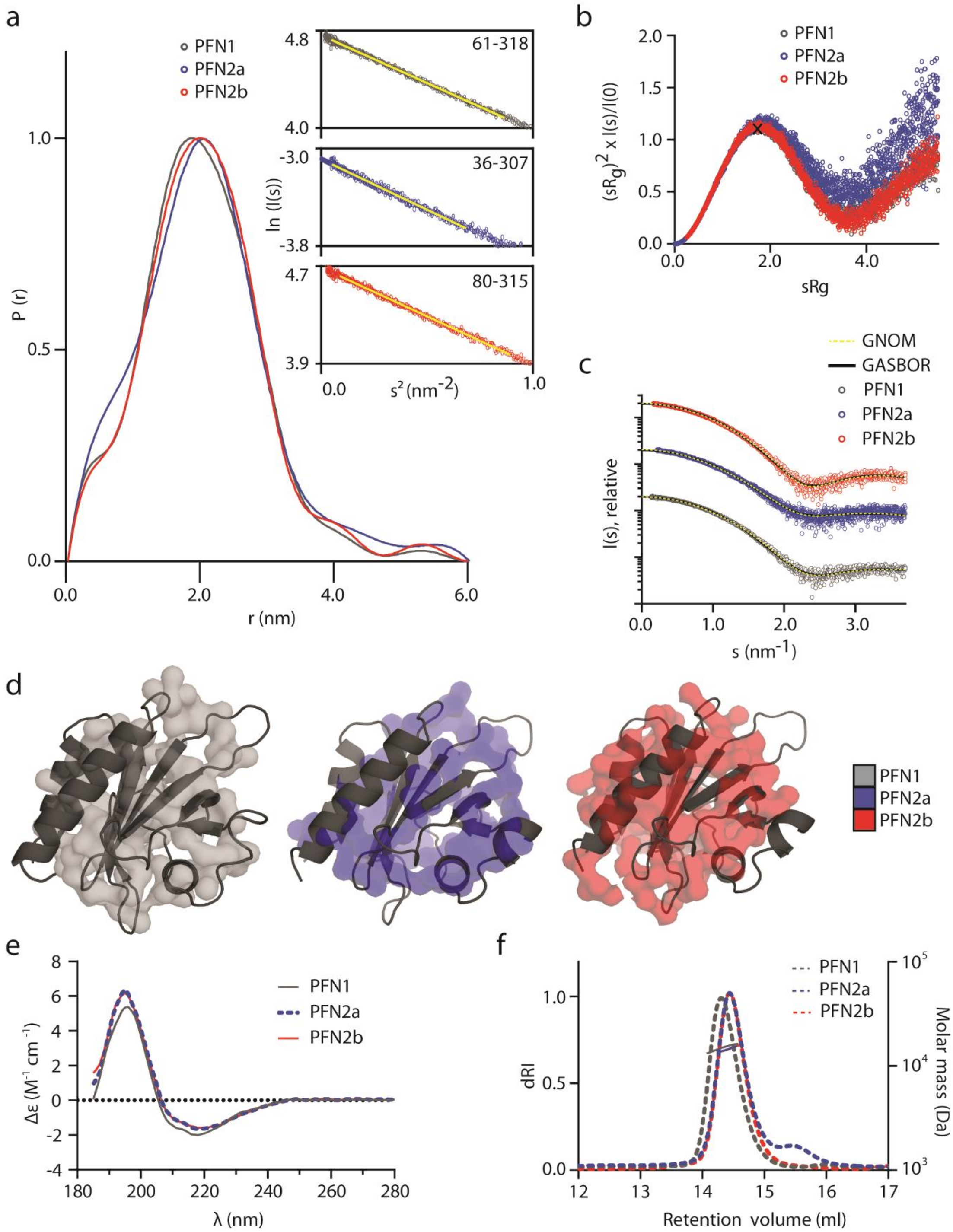

**Supplemental figure 4: SAXS, CD and MALS measurements of profilins.** a) Distance distribution of SAXS measurements of PFN1, PFN2a and PFN2b. Insert: Guinier plots of the same proteins. b) Dimensionless Kratky plot for PFN1, PFN2a, and PFN2b. The cross indicates the expected maximum for a rigid, spherical particle. c) Scattering profiles of PFN1, PFN2a and PFN2b. The data were shifted along the y-axis for clarity. The yellow, dotted line shows the GNOM fit, while the black line represents the GASBOR fit. d) GASBOR models of PFN1 (grey), PFN2a (blue) and PFN2b (red) overlaid with the crystal structures of each protein (surface, PFN1 PDB: 1FIK, PFN2 PDB: 1D1J). e) CD data for PFN1, PFN2a and PFN2b. f) SEC-MALS analysis of PFN1, PFN2a and PFN2b. Dotted lines show the differential refractive index recorded over the course of the SEC run (12 - 17 ml), while solid lines show the molecular mass of the particles in the peak.

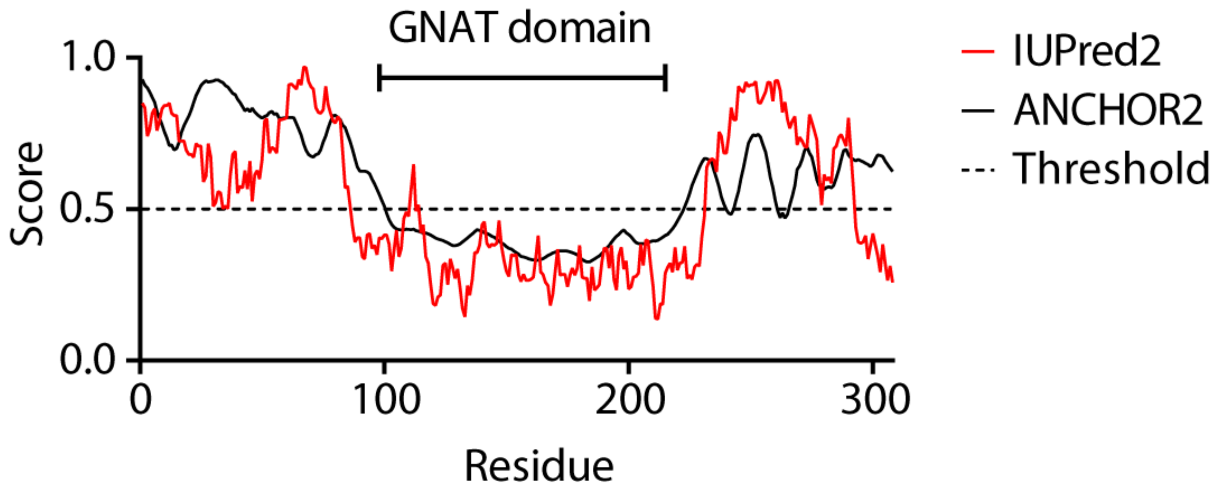

**Supplemental figure 5: Protein disorder prediction for NAA80, generated by IUPred2 and ANCHOR2 predictors.**

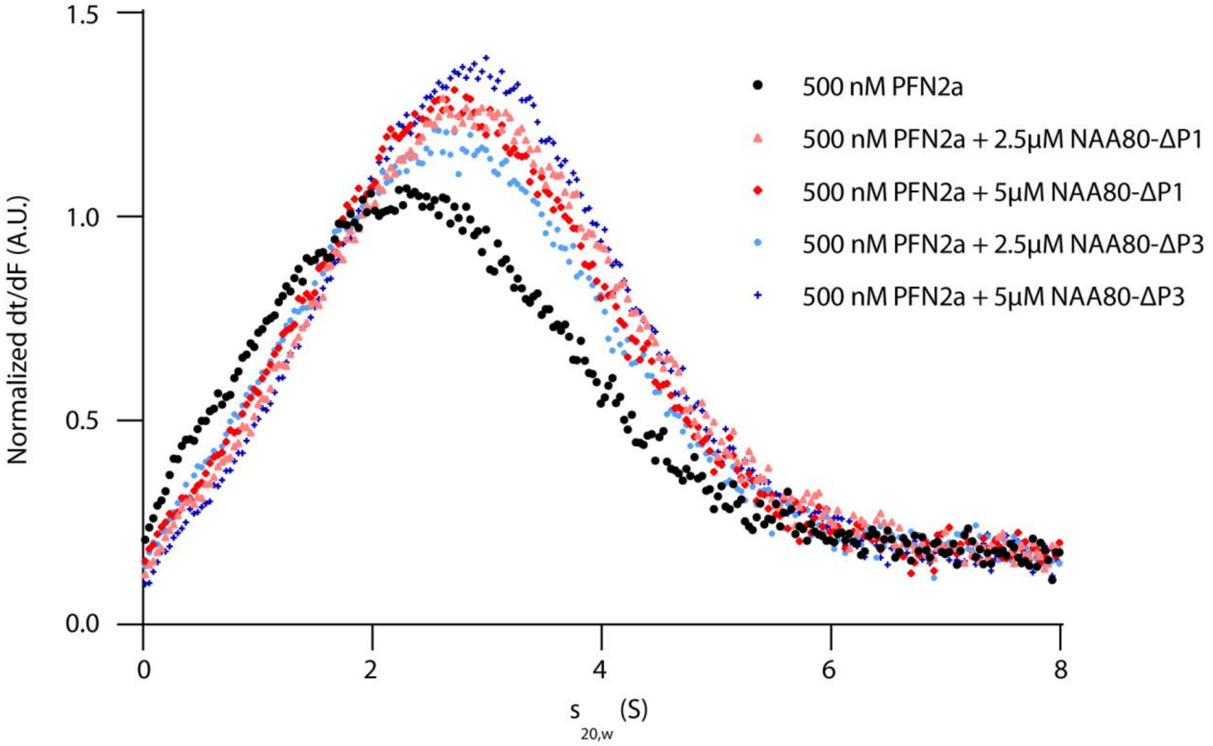

**Supplemental figure 6: Titration of NAA80-ΔP1 and NAA80-ΔP3 to PFN2a in AUC experiments.** AUC with labeled PFN2a and NAA80 deletion mutants, NAA80-ΔP1 or NAA80-ΔP3, in different concentrations.

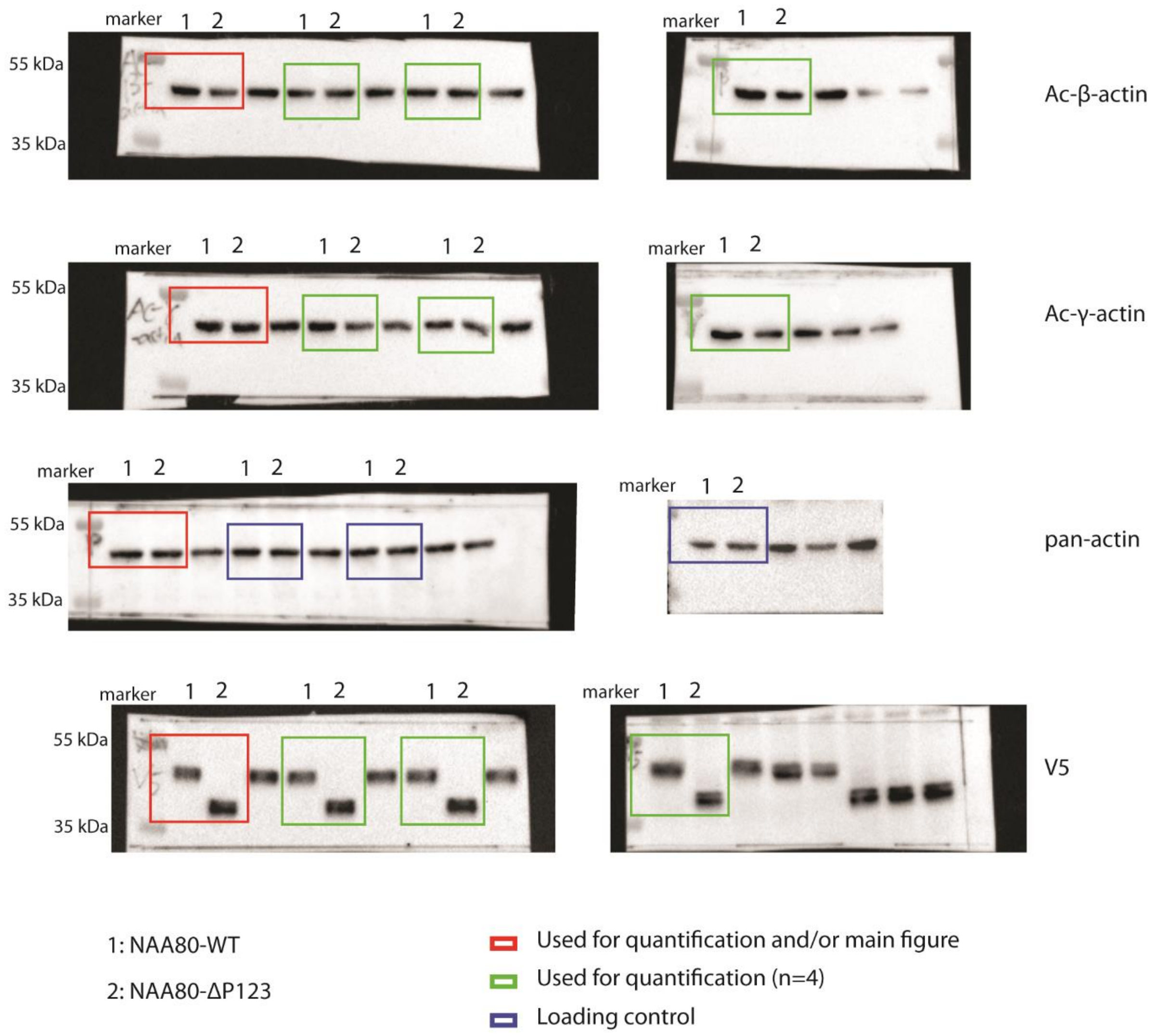

**Supplemental figure 7: Supplemental blots for the *in vivo* NAA80-WT and NAA80-ΔP123 actin reacetylation assay (Fig. 5c).** *NAA80*-KO cells were transfected with NAA80-WT or NAA80-ΔP123 (n = 4) for 14 hours and the lysates were probed with the indicated antibodies. Blots used in quantification (Ac-β-actin, Ac-γ-actin and V5) or as loading control (pan-actin) and included in the main figure (Fig. 5c) are outlined in red. Blots used only for quantification are outlined in green, and loading controls are outlined in blue.

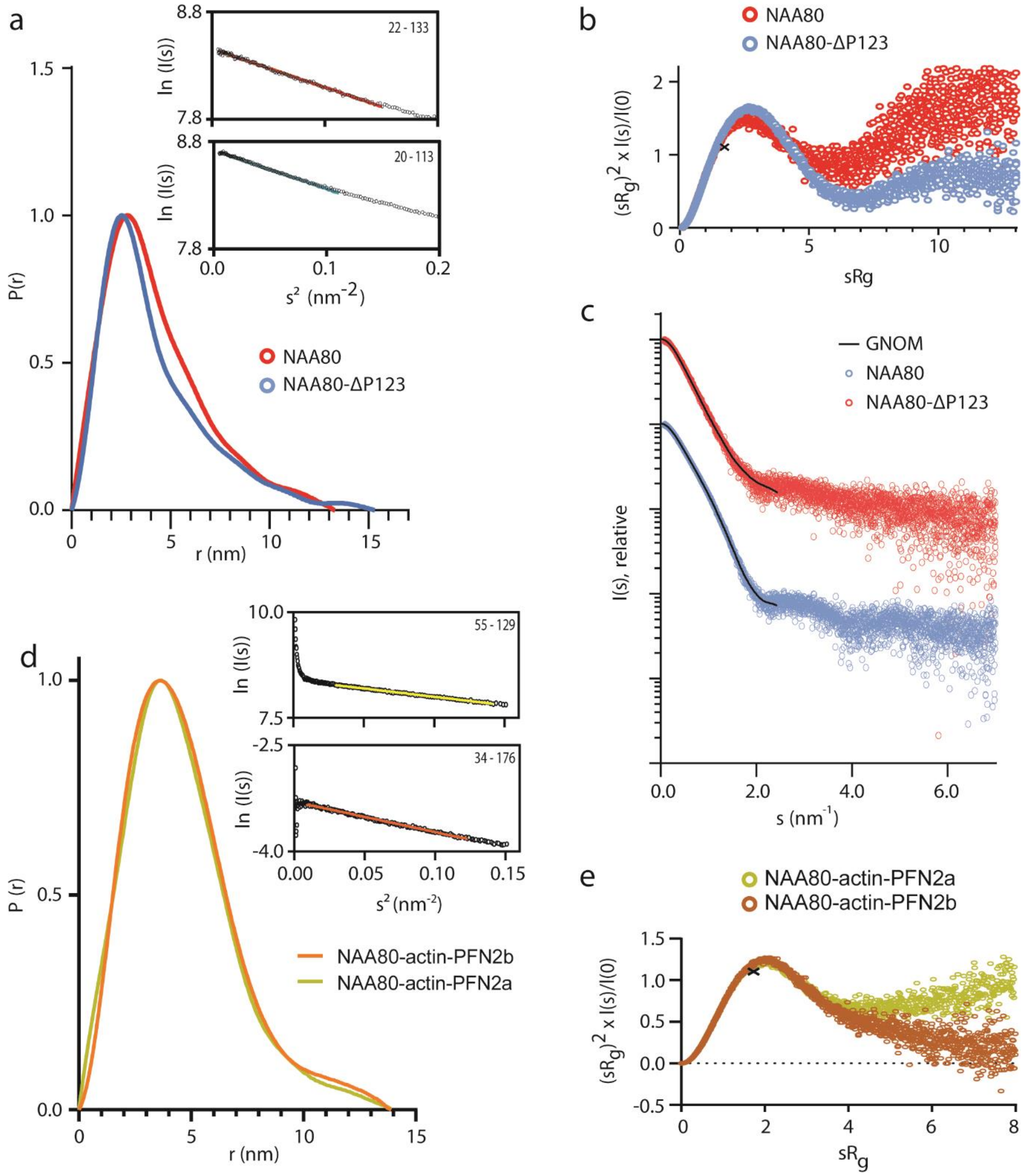

**Supplemental figure 8: SAXS measurements of NAA80, NAA80-ΔP123, and NAA80-actin-PFN2.** a) Distance distribution of SAXS measurements of NAA80 and NAA80-ΔP123. Inset: Guinier plots of the same proteins. b) Dimensionless Kratky plots for NAA80 (red) and NAA80-ΔP123 (blue). The cross indicates the expected maximum for a rigid, spherical particle. c) Scattering profiles of NAA80 (red) and NAA80-ΔP123 (blue). The data were shifted along the y-axis for clarity. The black line represents the GNOM fit (solid line). d) Distance distribution functions for NAA80-actin-PFN2a (yellow) and NAA80-actin-PFN2b (orange), with Guinier plots shown as inset. e) Normalized Kratky analysis plots for the same complexes. The cross indicates the expected maximum for a rigid, spherical particle.

**Supplementary tables 1 and 2: Excel sheets contain quantified proteins from LC/MS interactor screens in HeLa cells (Table S1) and HAP1 cells (Table S2).** Available for download online with this preprint. The mass spectrometry proteomics data and Tables S1-2 have additionally been deposited to the ProteomeXchange Consortium via the PRIDE partner repository with the dataset identifier PXD020188.

**Supplementary table 3: Thermal stability of NAA80 variants determined by DSF or CD (*)**

| **Protein** | $T_m$ **(°C)** | *s* **(°C)** |
|---|---|---|
| NAA80 | 41.75 | 0.07 |
| NAA80-ΔP1 | 42.65 | 0.25 |
| NAA80-ΔP2 | 42.15 | 0.25 |
| NAA80-ΔP3 | 41.28 | 0.07 |
| NAA80-ΔP123 | 41.06 | 0.17 |
| NAA80-PolyGS2* | 41.00 | 0.45 |

**Supplemental table 4: SAXS parameters for PFN1, PFN2a, and PFN2b**

| | | | |
|---|---|---|---|
| **Data collection** | | | |
| | **PFN1** | **PFN2a** | **PFN2b** |
| Instrument | P12, PETRA III, DESY, Hamburg | B21, Diamond Light Source, Harwell | P12, PETRA III, DESY, Hamburg |
| Wavelength (nm) | 0.124 | 0.100 | 0.124 |
| Angular range ($nm^{-1}$) | 0.04 – 7.3 | 0.04 – 3.7 | 0.04 – 7.3 |
| Temperature (°C) | 10 | 10 | 10 |
| SAXS mode | Batch | Batch | Batch |
| Exposure time (s) | 0.045 | 1 | 0.045 |
| Concentration (mg/ml) | 1.3 – 5.2 | 1.1 – 4.5 | 1.4 – 5.7 |
| **Software** | | | |
| Primary data reduction and processing | PRIMUS | | |
| Data validation and analysis | PRIMUS | | |
| *Ab initio* modelling | GASBOR | | |
| Graphical representation | PyMOL | | |
| **Structural data** | | | |
| I(0) (relative), from Guinier | 107.2 | 0.05 | 107.2 |
| $R_g$ (nm), from Guinier | 1.60 | 1.61 | 1.59 |
| s-range ($nm^{-1}$) used in Guinier | 0.22 – 0.93 | 0.23 – 0.83 | 0.30 – 0.95 |
| I(0) (relative), from P(r) | 107.2 | 0.05 | 107.2 |
| $R_g$ (nm), from P(r) | 1.60 | 1.61 | 1.60 |
| $D_{max}$ (nm), from P(r) | 6.0 | 6.0 | 6.0 |
| s-range ($nm^{-1}$) used in P(r) | 0.15 – 5.19 | 0.20 – 5.10 | 0.33 – 3.7 |
| **Molecular weight determination** | | | |
| $M_r$ (kDa), theoretical from sequence | 15.05 | 15.05 | 15.09 |
| $M_r$ (kDa), from I(0) | 14.30 | 14.60 | 14.30 |
| ***Ab initio* modelling** | | | |
| $\chi^2$ | 1.76 | 1.26 | 1.68 |

**Supplemental table 5: SAXS parameters for NAA80 and ternary complexes**

| **Data collection parameters** | | | | |
|---|---|---|---|---|
| | **NAA80** | **NAA80-ΔP123** | **NAA80-actin-PFN2a** | **NAA80-actin-PFN2b** |
| Instrument | P12, PETRA III, DESY, Hamburg | | B21, Diamond Light Source, Harwell | |
| Wavelength (nm) | 0.124 | | 0.1 | |
| Angular range ($nm^{-1}$) | 0.02 – 7.32 | | 0.03 – 4.4 | |
| SAXS mode | SEC-SAXS | | SEC-SAXS | |
| Injection volume (µl) | 100 | | 45 | |
| Exposure time (s/frame) | 1 | | 2.3 | |
| Number of frames | 3000 | | 1259 | |
| Temperature (°C) | 10 | | 10 | |
| **Software** | | | | |
| Data analysis software | ATSAS 2.8 | | | |
| Primary data reduction | CHROMIXS, PRIMUS | | | |
| Rigid body modelling | CORAL | | | |
| Ensemble optimization modelling | EOM | | | |
| Generation and fitting of theoretical scattering profiles | CRYSOL | | | |
| Graphical representation | PyMOL | | | |
| **Structural parameters** | | | | |
| I(0) (relative), from Guinier | 4695 | 6115 | 4425 | 0.037 |
| $R_g$ (nm), from Guinier | 3.27 | 3.25 | 3.44 | 3.58 |
| I(0) (relative), from P(r) | 4747 | 6181 | 4485 | 0.04 |
| $R_g$ (nm), from P(r) | 3.47 | 3.49 | 3.59 | 3.70 |
| $D_{max}$ (nm), from P(r) | 13.3 | 15.3 | 13.90 | 13.85 |
| Quality estimate | 0.71 | 0.66 | 0.79 | 0.75 |
| **Rigid body modelling** | | | | |
| Crystal structure | 6NAS:B | 6NAS:B | 6NAS | 6NAS |
| Flexible residues in NAA80 | 1 – 79, 217 – 303 | 1 – 79, 217 – 255 (256 – 302 missing) | 1 – 80, 221 – 258, 273 – 289 | 1 – 80, 221 – 258, 273 – 289 |
| $\chi^2$ | 3.25 | 23.1 | 1.45 | 1.16 |
| **Ensemble optimization method** | | | | |
| Crystal structure | 6NAS:B | 6NAS:B | | |
| Flexible residues in NAA80 | 1 – 79, 217 – 303 | 1 – 79, 217 – 255 (256 – 302 missing) | | |
| Number of conformers | 10 | 6 | | |
| $R_g$ (nm) | 3.35 | 3.20 | | |
| $D_{max}$ (nm) | 11.16 | 10.46 | | |
| $\chi^2$ | 0.95 | 1.45 | | |
| **Generation of theoretical scattering profile and SAXS profile fitting** | | | | |
| Crystal structure | 5WJD | | | |
| $R_g$ (nm) | 1.65 | | | |
| $D_{max}$ (nm) | 5.2 | | | |
| $\chi^2$ | 72.16 | | | |

**Supplemental table 6: List of primers, plasmids, antibodies and peptides used in this study.**

| Primer ID | Sequence | Plasmid product/Usage |
|---|---|---|
| oTA22: NAA80 KozF | caacatgcaagagctgactc | pTA166 |
| oTA792: NAA80 w STOP rv | tcagatgtctttttccatccagaatatg | pTA166 |
| oTA732: NAA80 fw | Atgcaagagctgactctgagc | pTA538 |
| oTA311: NAA80 w STOP | Catgaattccattcagatgtctttttccatcc | pTA538 |
| oTA812: NAA80 M23L fw | tacacacccggctggagctgat | pTA587, introducing M23L |
| oTA813: NAA80 M23L rv | Gggtctagtgtaggggtc | pTA587, introducing M23L |
| oTA864: pcDNA4 Xpress to V5 fw | ctcctcggtctcgattctacggtacaggccccttatgcaag | pTA609, replacing N-terminal Xpress tag with V5 tag |
| oTA865: pcDNA4 Xpress to V5 rv | agggttagggataggcttaccccgacccatttgctgtcc | pTA609, replacing N-terminal Xpress tag with V5 tag |
| oTA904: NAA80□238-247 fw | aacctgactgcccaagctg | pTA646, pTA777, pTA780 |
| oTA905: NAA80□238-247 rv | Ggctgtggggaaggcatt | pTA645, pTA777 |
| oTA906: NAA80□277-284 fw | Tcaaaaagcctgctggag | pTA779 |
| oTA907: NAA80□277-284 rv | Tgagatggtcaggcactc | pTA779, pTA780 |
| oTA917: NAA80□260-270 fw | gagtgcctgaccatctcaa | pTA778 |
| oTA918: NAA80□260-270 rv | cttgggaccccttggggca | pTA778 |
| oTA1146: NAA80 polyGS2 fw | ggctctggcctatctgagtgcctgaccatctcac | pTA858 |
| **oTA1147: NAA80 polyGS2 rv** | **agagcccaagctgcctcccttgggaccccttgg** | **pTA858** |

-

| Plasmid ID | Description | Expression | Protein |
|---|---|---|---|
| pTA12 | pcDNA3.1-lacZ-V5 | Mammalian | β-galactosidase-V5 |
| pTA166 | pcDNA3.1-NAA80-V5 | Mammalian | NAA80-V5 |
| pTA538 | pcDNA4-Xpress-NAA80 | Mammalian | Xpress-NAA80 |
| pTA587 | pcDNA4-Xpress-NAA80-M23L | Mammalian | Xpress-NAA80 (M23L) |
| pTA598 | pTYB11-profilin-1 | Bacterial | PFN1 |
| pTA599 | pTYB11-profilin-2a | Bacterial | PFN2a |
| pTA600 | pTYB11-profilin-2b | Bacterial | PFN2b |
| pTA609 | pcDNA4-V5-NAA80 | Mammalian | V5-NAA80 (M23L) |
| pTA646 | pcDNA4-V5-NAA80-Δ238-284 | Mammalian | V5-NAA80 (M23L) Δ238-284 (ΔP123) |
| pTA631 | pCold1-Gelsolin C'term half (G4-G6) | Bacterial | Gelsolin |
| pTA858 | pcDNA4-V5-NAA80-polyGS2 | Mammalian | V5-NAA80 (M23L) polyGS2 |
| pTA751 | pTYB12-NAA80 | Bacterial | NAA80 |
| pTA777 | pTYB12-NAA80-Δ238-247 | Bacterial | NAA80-Δ238-247 (ΔP1) |

| PRIMARY ANTIBODIES | | | | |
|---|---|---|---|---|
| **Antibody** | **Cat. Nr** | **Company** | **Animal** | **Dilution** |
| Pan-actin | AAN01 | Cytoskeleton | Rabbit | 1:2000 |
| Pan-actin | Ab14128 | Abcam | Mouse | 1:2000 |
| Ac-β-actin | ab6276 | Abcam | Mouse | 1:3000 |
| Ac-γ-actin | ab123034 | Abcam | Mouse | 1:5000 |
| Profilin-1 (PFN1) | ab50667 | Abcam | Rabbit | 1:2000 |
| Profilin-2 (PFN2) | SC-100955 | Santa Cruz | Mouse | 1:1000 - 1:2000 |
| V5 | R960CUS | Invitrogen | Mouse | 1:1000 - 1:5000 |
| α-Vinculin | ab129002 | Abcam | Rabbit | 1:1000 - 1:2000 |
| NAA80 | custom | Biogenes | Rabbit | 1:200 |

| SECONDARY ANTIBODIES AND STAINING REAGENTS | | | | | |
|---|---|---|---|---|---|
| **Antigen** | **Cat. Nr** | **Company** | **Animal** | **Dilution** | **Usage** |
| ECL Anti-mouse IgG-HRP | NA931 | Amersham | Goat | 1:3000-1:20000 | Western blotting |
| ECL Anti-rabbit IgG-HRP | NA934 | Amersham | Goat | 1:3000-1:20000 | Western blotting |

| Name | Oligopeptide sequence | NAT | Derived from | UniProt |
|---|---|---|---|---|
| DDDI | [H] **DDDIAAL** RWGRPVGRRRRPVRVYP [OH] | H | β-cyto-actin | P60709 |
| EEEI | [H] **EEEIAAL** RWGRPVGRRRRPVRVYP [OH] | H | γ-cyto-actin | P63261 |
